# Theory of sarcomere assembly inferred from sequential ordering of myofibril components

**DOI:** 10.1101/2023.08.01.551279

**Authors:** Francine Kolley, Clara Sidor, Benoit Dehapiot, Frank Schnorrer, Benjamin M. Friedrich

## Abstract

Myofibrils in striated muscle cells are chains of regular cytoskeletal units termed sarcomeres, whose contractions drive voluntary movements of animals. Despite the well characterized order of the sarcomere components in mature sarcomeres, which explains the sarcomere contraction mechanism, the mechanism of molecular ordering during sarcomere assembly remains debated. Here, we put forward a theoretical framework for the self-assembly of sarcomeres. This theory is based on measurements of the sequential ordering of sarcomere components in developing *Drosophila* flight muscles, identified by applying a novel tracking-free algorithm: myosin, *α*-actinin and the titin homologue Sallimus form periodic patterns before actin. Based on these results, we propose that myosin, Sallimus, and sarcomere Z-disc proteins including *α*-actinin dynamically bind and unbind to an unordered bundle of actin filaments to establish an initial periodic pattern. As a consequence, periodicity of actin filaments is only established later. Our model proposes that non-local interactions between spatially extended myosin and titin/Sallimus containing complexes, and possibly tension-dependent feedback mediated by an *α*-actinin catch-bond, drive this ordering process. We probe this hypothesis using mathematical models and derive predictive conditions for sarcomere pattern formation, guiding future experimental analysis.

## Introduction

Striated muscle as well as heart muscle cells contain acto-myosin bundles of almost crystalline regularity, termed myofibrils, which span the entire multi-nucleated muscle cell with a length of up to several millimeters [1]. Activation of myosin activity results in the contraction of myofibrils [2, 3], which powers all voluntary movements in humans and animals, as well as heartbeat. Each myofibril is a periodic chain of stereotypic units, termed sarcomeres, with a well-defined, micrometer-range length. Each sarcomere is bordered by two *α*-actinin-rich Z-discs, which cross-link parallel, polarity-sorted actin filaments at their plus-end (barbed end), see also Fig. 1, right. Actin filament minus-ends face towards central, bipolar myosin filaments, cross-linked by myomesin (or Obscurin in *Drosophila*). Importantly, the giant protein titin (Sallimus in *Drosophila*), anchored at the Z-disc, stably links actin (thin filaments) and bipolar myosin filaments (thick filaments) together by extending from the Z-disc to the center of the sarcomere in mammals [4] or to the beginning of myosin filaments in flight muscles [5]. Defects in sarcomere assembly result in severe myopathies [6, 7].

**FIG. 1.**
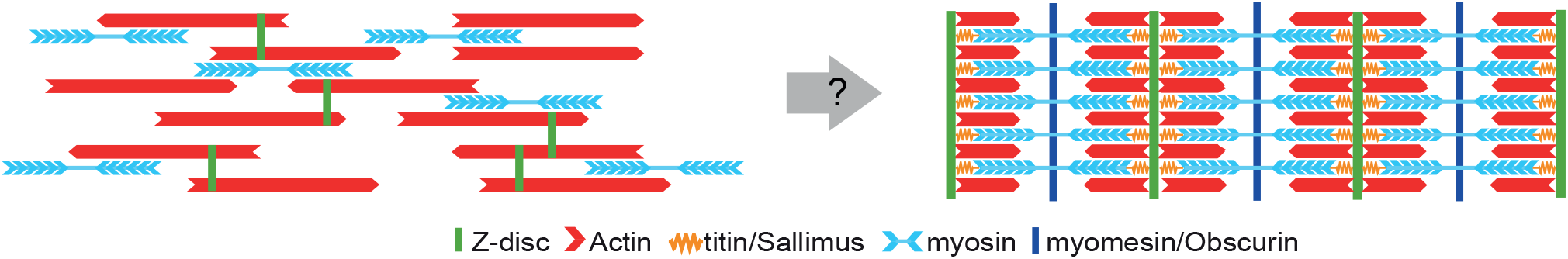
Myofibrillogenesis represents a pattern formation problem: How do initially stress-fiber like bundles of parallel actin filaments (red) with homogeneous distribution of myosin (blue) and Z-disc proteins (green) re-arrange into periodic sarcomeric patterns? In mature myofibrils, actin filaments form polarity-sorted domains, with their plus-ends crosslinked at the Z-disc rich in *α*-actinin, which marks the boundary of the sarcomere. Bipolar myosin filaments in the middle of each sarcomere are crosslinked by myomesin/Obscurin (dark blue) and anchored to the Z-discs by titin/Sallimus (yellow).

Despite many years of research on sarcomere and myofibril development, we still do not understand the physical mechanisms that drive this self-assembly process. In particular, it remains open why mechanical tension is essential for sarcomere formation *in vivo* and *in vitro* [8–12], prompting for physical descriptions. Sanger and colleagues proposed in the premyofibril hypothesis that non-muscle myosin and Z-disc proteins establish early periodic patterns, with non-muscle myosin being replaced by muscle-myosin later [13]. This model may apply in some muscles; however, the periodicity of non-muscle myosin is not always clearly visible [10, 14]. Irrespective of whether early myofibrils comprise non-muscle or only muscle-specific myosin filaments, the key question is the following: how do the first periodic patterns form?

Holtzer and colleagues observed small I-Z-I bodies comprising Z-protein aggregates, as well as free-floating stacks of bipolar myosin filaments in atypical myogenic cells [15], leading to the proposal that these “building blocks” may become stitched together at later stages to form sarcomeres. Latent protein complexes were also proposed as precursors for *Drosophila* larval muscles [16], or chicken cardiomyocytes [17]. However, it is not clear how such large supra-molecular complexes could move in the crowded environment of a muscle cell.

Previous mathematical models commonly assumed that periodic patterns of polarity sorted actin filaments form simultaneously with periodic patterns of myosin and Z-disc proteins [18–20]. Zemel et al. showed how a hypothetical second, minus-end directed molecular motor could establish periodic cytoskeletal patterns [18, 21]. Yoshinaga et al. showed how a generic coupling between actin polarity and stress fields result in the same pattern [22]. Finally, Friedrich et al. proposed actin polymerization forces as a physical mechanism for the polarity sorting of actin filaments into a periodic pattern [20]. None of these theoretical mechanisms have been confirmed by experimental data so far, and some mechanisms pose open questions, for example, actin polymerization forces may be too weak to drive filament sorting [23, 24]. Furthermore, titin, which was shown to be indispensable for myofibril assembly [25–27], and proposed to directly affect sarcomere length [4, 28–30], had not been included in these previous models. More generally, theoretical descriptions of cytoskeletal pattern formation in pure actomyosin systems predict localization of myosin filaments near the plus-end of actin filaments [31, 32], opposite to their localization in mature sarcomeres.

Here, quantifications of periodic patterns in assembling myofibrils of *Drosophila* muscles, lead us to propose a new model: our data suggest that myosin and Z-disc proteins establish the first periodic patterns, while actin forms polarity-sorted pattern only later. We postulate that myosin and Z-disc proteins bind and unbind to and from a bundle of actin filaments, which are aligned parallel or anti-parallel, and thus are not yet polarity sorted. Reciprocal interactions between myosin and Z-disc proteins, possibly mediated indirectly through titin, favor the formation of periodic sarcomeric patterns. Only subsequently, will actin become polarity-sorted, possibly by preferential nucleation of actin filaments at emergent periodically positioned Z-bodies, and gradual depolymerization of ectopic actin filaments.

We additionally introduce a second, modified model based on mechanical tension generated by myosin motor activity. We show that the proposed catch-bond behavior of the Z-disc protein *α*-actinin [33–35] is sufficient to drive the formation of periodic patterns in simulations, thus providing a mechanistic underpinning for the idea that effective elastic interactions between myosin bipolar filaments may drive sarcomeric pattern formation [36].

## RESULTS

### Sequential emergence of periodic patterns during myofibrillogenesis

In *Drosophila melanogaster*, myofibrillogenesis is a multi-stage process, in which sarcomere assembly starts after 22 h after pupa formation (APF). Unstriated acto-myosin bundles with nematically ordered actin filaments are present at 22 h, and sub-sequentially mature into myofibrils by 32 h APF [8, 37]. To gain insight into the gradual establishment of periodic sarcomeric patterns in developing myofibrils, we obtained multichannel fluorescence z-stack images of dorsal longitudinal flight muscles (DLMs4) from *Drosophila melanogaster* stained for *α*-actinin, actin, myosin and the titin homologue Sallimus (Sls) at selected time points, see Fig. 2A.

**FIG. 2.**
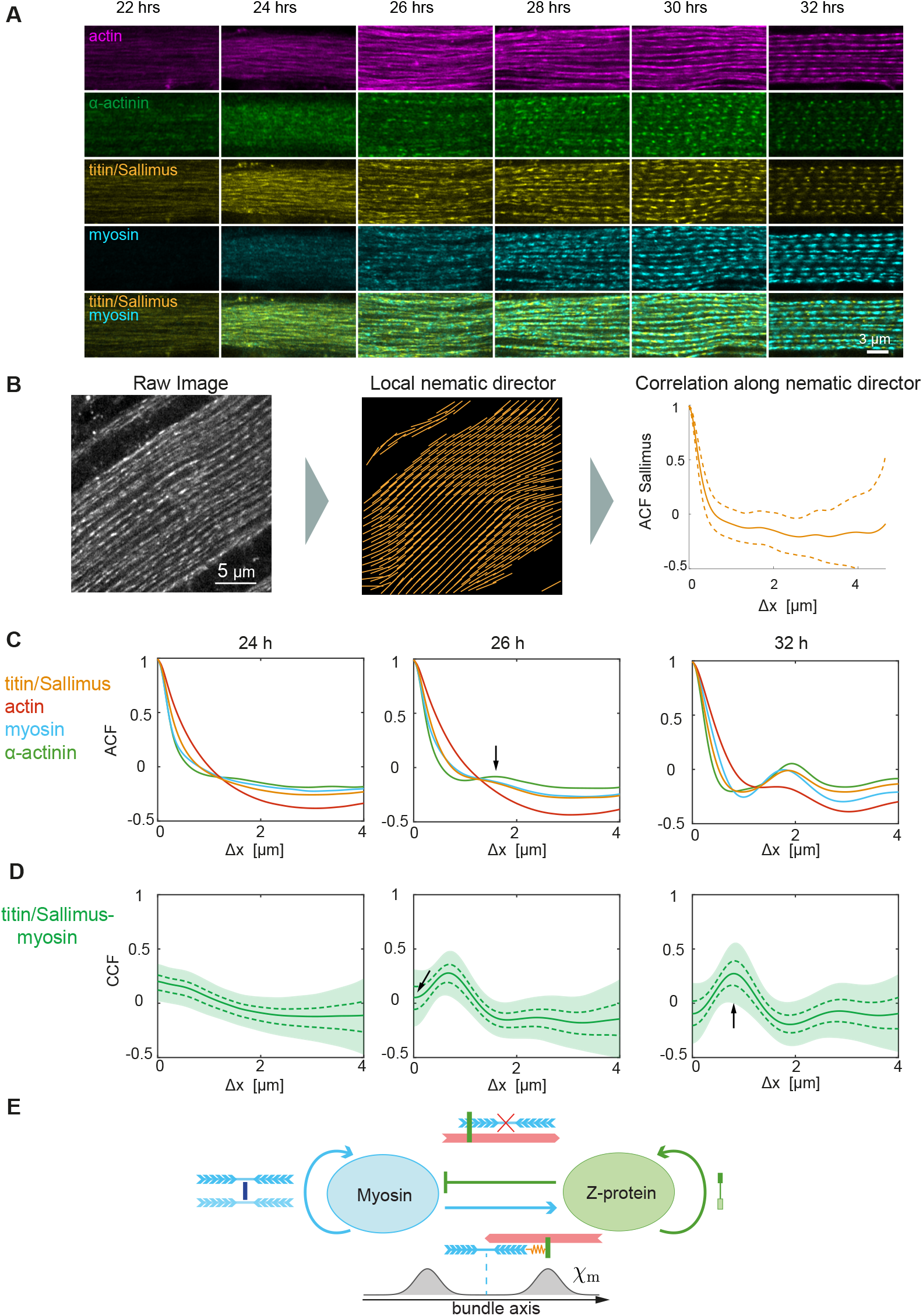
**A**. Multi-channel images of *Drosophila* flight muscle at 22 h - 32 h after pupae formation (APF) with actin (magenta), myosin (blue), titin/Sallimus (yellow) and myosin-Sallimus merge. Scale bar 5 *μ*m. **B**. Tracking-free algorithm to compute correlation functions along local bundle axis. After pre-processing, a field of local nematic directors is determined (middle). Linescans along these nematic directions enable to compute mean correlation functions (right). **C**. Auto-correlation functions (ACF) for actin (red), myosin (blue), Sallimus (yellow), *α*-actinin (green) at selected time-points. A monotonically decreasing ACF indicates a random distribution of proteins, while a peak (arrow) reveals periodic patterns with wavelength given by peak position. **D**. Cross-correlation function (CCF) between myosin and Sallimus (green, mean *±* s.e.m. (dashed), *±*s.d. (shaded)). A positive peak at non-zero Δ*x* indicates a localization of the two proteins at a characteristic distance, while a negative dip at Δ*x* = 0 indicates a local exclusion. **E**. Proposed model of non-local interactions between myosin and Z-disc proteins driving pattern formation. Lateral interactions between bipolar myosin filaments and crosslinkers favor autocatalytic attachment. Lateral interactions between bipolar myosin filaments favor autocatalytic binding of myosin to the actin bundle. Likewise, interactions between Z-disc proteins favor autocatalytic binding of Z-disc proteins. Steric repulsion by Z-disc proteins impedes myosin binding to the actin bundle, causing a negative feedback. Finally, myosin recruits Sallimus, which recruits Z-disc proteins such as *α*-actinin. This positive feedback is non-local, here modeled by an interaction kernel *χ*_*m*_ with mean interaction distance *l*_*m*_*/*2 + *l*_*s*_ set by the length *l*_*m*_ of myosin and *l*_*s*_ of the giant protein Sallimus.

No periodic patterns are observable by eye at 22 h and 24 h APF. At 26 h, first periodic patterns with alternating localization of *α*-actinin, Sallimus and myosin start to emerge. Actin, however, does not yet display obvious periodic patterns at 26 h. Over time, periodic patterns become increasingly more pronounced, and are clearly visible for all four sarcomere components at 32 h (Fig. 2A). Thus, sarcomere proteins assemble into periodic patterns from 26 h to 32 h APF.

### Microscopy methods

*Drosophila melanogaster* pupae of the *Mhc* (*weeP26*) strain, which expresses a myosin heavy chain (Mhc) endogenously tagged with a Green Fluorescent Protein (GFP), were grown at 27°C for different durations from 22 h up to 32 h APF, which corresponds to the time frame of sarcomere assembly in the indirect flight muscles at this temperature [8, 38]. Actin was labeled with phalloidin Alexa-568, Sallimus with the fluorescent nanobody Sls-Nano2-DyLight405 and *α*-actinin with Actn-Nano62-Atto647 [27], see Supplemental Material (SM) for additional details on experimental methods.

### A robust, tracking-free algorithm to compute correlations functions

To quantify the order of individual proteins in these micrographs, we developed a tracking-free image-analysis algorithm to compute correlation functions of protein intensities along myofibrils, Fig. 2B see also Fig. S1. This algorithm does not track individual myofibrils, but uses the estimated local direction of myofibrils. Even manual tracking of individual myofibrils is impossible at early stages because myofibrils have not formed yet. Specifically, after pre-processing, a steerable filter was used to determine the local nematic direction of the Sallimus channel in non-overlapping regions-of-interest (ROI) (Fig. 2B). Next, intensity profiles *I*(*x*) were computed for each channel from 5 *μ*m line-scans along these local nematic directions for each ROI. Auto-correlation functions (ACFs) and cross-correlation functions (CCFs) were then computed from these intensity profiles, and averaged over all ROIs, as 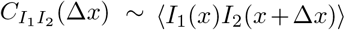, see Supplemental Material for details.

### Correlation functions reveal temporal order of sarcomeric components and suggest interactions

Fig. 2C shows computed auto-correlation functions for *α*-actinin, myosin, Sallimus, and actin at three selected time points (for additional time points, see Supplemental Material). A Fourier peak at a position Δ*x* in the ACF reveals periodic patterns with characteristic periodicity Δ*x*, even if patterns are noisy. More precisely, a Fourier peak at position Δ*x* of amplitude *A* indicates periodic order with periodicity *L* = Δ*x* [1 + tan^−1^(ln(*A*)*/π*)*/*(2*π*)]^−1^ ≥ Δ*x*, which implies a small correction for noisy patterns, see SM for details. In contrast, a random localization of a protein without any periodic pattern is reflected by a monotonically decreasing ACF. We applied this method to the developing flight muscle images. At 22 h and 24 h APF, the ACFs display no evidence for periodic patterns for any of the four proteins (actin, myosin, Sls, *α*-actinin) tested (Fig. 2C). A first indication for periodic patterns *α*-actinin, Sallimus and myosin is found at 26 h APF as a small Fourier peak located at Δ*x* ≈ 1.5 *μ*m (Fig. 2C). The slightly larger peak of *α*-actinin suggests an initially more disperse distribution of myosin and Sallimus. The amplitude of these Fourier peaks increases with time, reflecting the formation of increasingly regular patterns. The position of the Fourier peaks shift to Δ*x* ≈ 2.1 *μ*m at 32 h APF, confirming the known increase of sarcomere size during the beginning of *Drosophila* myofibrillogenesis [39].

This length increase could result from myosin filaments growing in length [15], or mechanical tension that stretches sarcomeres, which was found to be high at this stage [40]. In contrast, the ACF for actin is still monotonic decreasing up to 30 h APF, and only shows a Fourier peak at 32 h APF (Fig. 2C). This observation strongly suggests that myosin, *α*-actinin and Sallimus form periodic patterns first, while actin follows 6 hours later.

In addition to ACFs, we can compute cross-correlation functions (CCF) between different channels. As an example, Fig. 1D displays the CCF between Sallimus and myosin: at 26 h APF, a negative peak at Δ*x* = 0 indicates an anti-correlation between Sallimus and myosin filaments. This anti-correlation is a strong indication for a local exclusion between these molecular players, consistent with the hypothesis of a direct or indirect negative interaction.

To assess possible spatial variations in myofibril maturation, we selected the 5% of ROIs in fluorescence images with most progressed myofibrillogenesis (according to the height of the Fourier peak in the ACF for Sallimus), and re-computed correlation functions, which gave almost identical results with a time shift of 2 h at most, showing the high degree of synchronization in pattern assembly, see Fig. S4 in the SM.

In conclusion, our experimental data show that myosin and the titin-homologue Sallimus, which links Z-disc components to myosin, assemble into a periodic pattern before actin does. This assembly is largely homogeneous throughout the large muscle cells. At 24 h APF, small irregularities are already observable before a global periodic pattern emerges at 26 h APF.

### A new mechanistic hypothesis

Based on previous work [8, 37, 39] and our observations from early stages of myofibrillogenesis in *Drosophila* flight muscle, we put forward two putative mechanisms that suggest how the selforganized formation of sarcomeres may happen. In both proposed mechanisms, myofibrillogenesis starts from initially unstriated bundles of nematically aligned actin filaments that are not polarity-sorted yet, but are nematically ordered, i.e., aligned parallel or anti-parallel to the bundle axis, resembling stress fibers in connective tissue cells. Myosin, Z-disc proteins and Sallimus dynamically bind and unbind from these bundles, interacting in such a manner that regular patterns with alternating localization and correct periodicity emerge. Periodic actin patterns will become established later, possibly driven by continuous actin turnover, preferential nucleation at nascent Z-discs (or the M-band [41]), and depolymerization of ectopic actin filaments, see Discussion section.

We assume that bipolar myosin filaments bound to the initial actin scaffold will recruit more myosin, representing autocatalytic attachment or *in situ* polymerization of myosin. This assumption is warranted by the experimental observation that bipolar myosin filaments form ordered stacks (and that existing stacks can “catch” new myosin filaments), as observed in non-muscle cells [42], human osteosarcoma cells [43], and in atypical myogenic cells [15]. Moreover, aligned myosin filaments become crosslinked by M-band proteins (e.g., myomesin or Obscurin in the fly) [44].

Spontaneous aggregation was described also for Z-disc proteins, characterized by Z-disc protein punctae that fuse into larger nascent Z-discs [45]. Similar Z-disc protein aggregates had been observed previously in Taxol-treated cells [15]. Together, these observations strongly suggest autocatalytic aggregation of Z-disc proteins.

Fig. 2E summarizes these putative molecular interactions between myosin and Z-disc proteins bound to actin. In addition to autocatalytic attachment of both myosin and Z-disc proteins, we assume that Z-disc proteins locally inhibit myosin binding due to steric repulsion by the extended titin/Sallimus protein, reflecting a competition for actin binding sites between Z-disc proteins and myosin filaments. This creates a negative feedback loop (Fig. 2E). Complementary, we propose that myosin enhances the binding of Z-disc proteins, reflecting the recruitment of Sallimus by myosin, which then recruits Z-disc proteins such as *α*-actinin at its N-terminus. For this, we assume that Sallimus attains its extended configuration already at early stages of myofibrillogenesis [5].

Importantly, interactions between the spatially extended molecules define non-local interactions spanning over a distance set by the lengths of myosin filaments and Sallimus molecules (mathematically modeled as a double-peak interaction kernel with its half-length set by the half-length of myosin filaments *l*_*m*_*/*2 and the length of Sallimus *l*_*s*_, sketched in Fig. 2E). Experimental data suggest that giant proteins such as titin act as molecular rulers that, directly or indirectly, orchestrate sarcomeric length control [4, 28–30]. In particular, expression of shorter titin isoforms caused a decrease of mature sarcomere length [4, 29, 30]. In our mathematical model, the interaction length of non-local interactions turned out to be an important determinant of sarcomere length.

Next, we formulate the model shown in Fig. 2E as a minimal mathematical model referred to as model I, and demonstrate its capability to establish regular sarcomeric patterns using agent-based simulations.

### Sarcomeric pattern formation by non-local interactions

We translate the minimal mathematical model I, which assumes molecular interactions between myosin and Z-proteins as proposed in Fig. 2E, into a minimal mathematical model. The model couples the concentration *m*(*x*) of bound myosin filaments and *z*(*x*) of bound Z-disc proteins as function of the bundle axis coordinate *x* through binding and unbinding rates that depend on the number of molecules already bound, while accounting for small random movements of bound myosin and Z-disc proteins along the fiber axis due to random forces, modeled as apparent diffusion with effective diffusion coefficient *D*

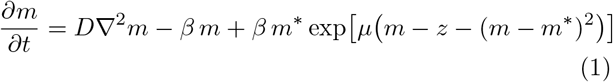

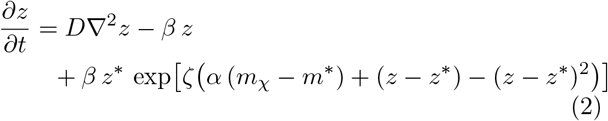

For simplicity, we assume equal effective diffusion coefficients *D* = *D*_*m*_ = *D*_*z*_, equal unbinding rates *β* = *β*_*m*_ = *β*_*z*_ = *β*, as well as equal steady-state concentrations *m*^*^ = *z*^*^ for myosin and Z-disc proteins, see Supplemental Material for a generalization. As a technical point, the concentration *m*(*x*) refers to the midpoints of extended myosin filaments.

The binding rate of myosin in Eq. (1) is modulated from the base rate *β* depending on the concentrations *m*(*x*) and *z*(*x*) of myosin and Z-disc proteins already bound to account for the molecular interactions reviewed in Fig. 2E. The proposed autocatalytic binding of myosin is encompassed by the exponential factor exp(*μm*), while the factor exp(−*μz*) represents negative feedback of bound Z-disc protein on myosin recruitment due to steric interactions. The quadratic term in the exponential captures saturation effects that limit deviations from the steady-state concentration *m*^*^.

Eq. (2) for *z*(*x*) is similar, with the important difference that the factor exp(*ζαm*_*χ*_) describes a non-local interaction between extended myosin filaments and Z-disc proteins, mediated by the giant protein Sallimus. Mathematically, *m*_*χ*_ = *m* * *χ*_m_ in Eq. (2) is a convolution of the local concentration of bound myosin *m*(*x*) and a non-local interaction kernel *χ*_m_(*x*), taken as a double-Gaussian kernel with peaks of width *σ* located at a distance ±*l*_*χ*_, see also Fig. 1E. The interaction length *l*_*χ*_ = *l*_m_*/*2 + *l*_s_ is set by the lengths *l*_m_ and *l*_s_ of myosin filaments and Sallimus. The parameter *α* allows to tune the strength of this non-local interaction.

A linear stability analysis of Eqs. (1-2) reveals parameter regimes, for which the homogeneous steady state with *m*(*x*) ≡ *m*^*^ and *z*(*x*) ≡ *z*^*^ is unstable, and small spatial inhomogeneities become amplified. Indeed, numerical integration of Eqs. (1-2) results in regular periodic patterns with alternating peaks of bound myosin and Z-disc proteins, see Fig. 3B for an example and Supplemental Material for details.

**FIG. 3.**
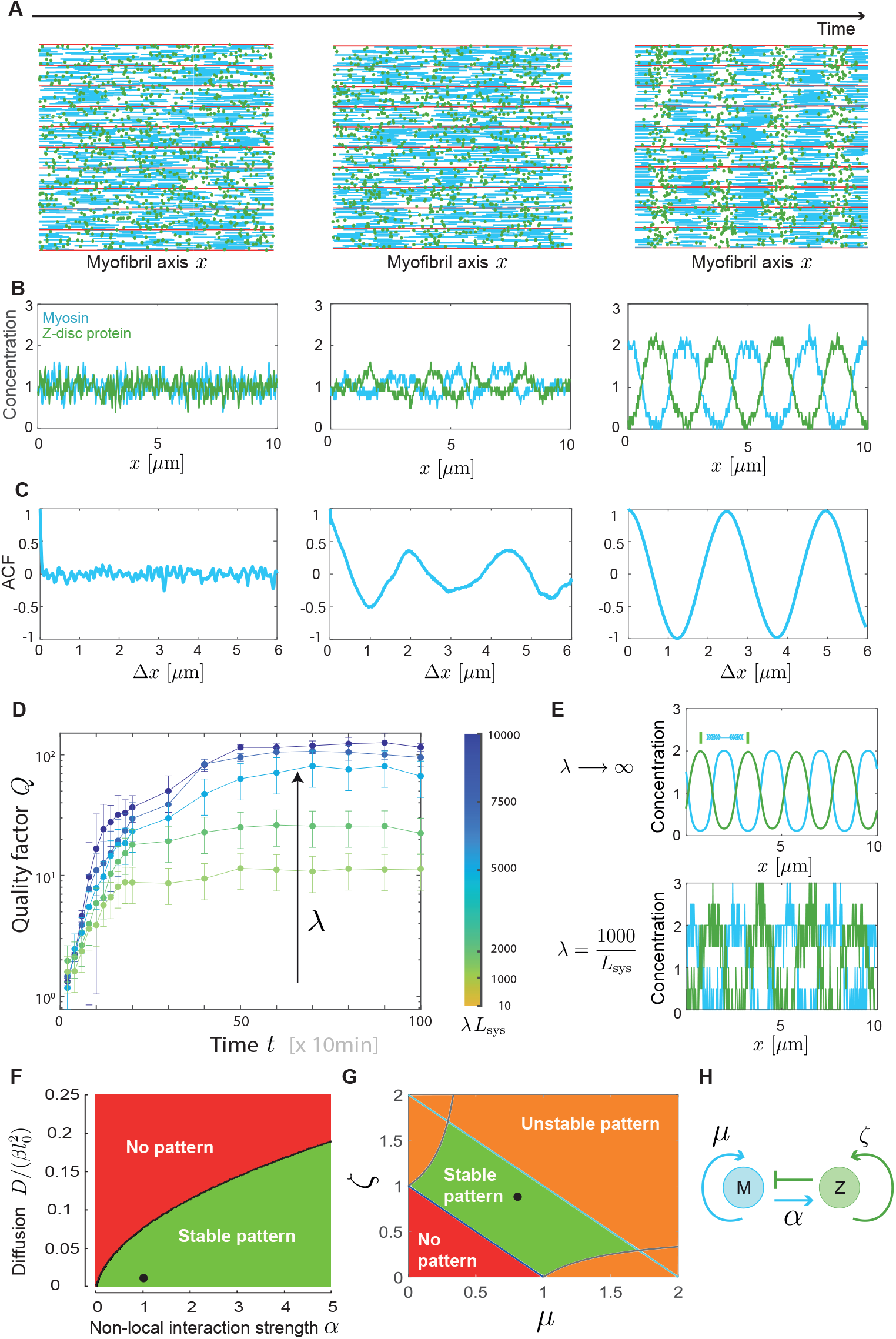
Pattern formation by non-local interactions. **A**. Agent-based simulations of mathematical model I proposed in Fig. 1B, simulated in one space dimension, visualized in two space dimensions, for different simulation times (*t* = 0, 2, 100) with myosin (blue) and Z-disc protein (green) bound to a scaffold of parallel, infinitely long actin filaments (shown schematically, red). **B**. Corresponding concentration profiles at *t* = 0, 2, 100 for myosin (blue) and Z-disc protein (green). **C**. Auto-correlation functions (ACF) at *t* = 0, 2, 100 for myosin concentration. The ACF for *t* = 0 fluctuates around zero, reflecting the initially random distribution of myosin. The ACF for *t* = 2 exhibits a Fourier peak with amplitude *A*, indicative of a periodic pattern. **D**. Quality factor *Q* = −*π/* ln(*A*) of periodic patterns determined from the amplitude *A* of the Fourier peak in the ACF of myosin concentration as function of time *t* for different values of total myosin density *λ* (mean*±*s.e.m., *n* = 10 simulation runs). **E**. Example concentration profile for different values of *λ*, corresponding to different steady-state values of the quality factor *Q*. A mean-field model was used for the limit *λ* → ∞. **F**. Phase diagram showing regimes of stable patten formation as function of the strength *α* of non-local interactions between myosin and Z-disc proteins, and the diffusion coefficient *D* of bound myosin filaments. **G**. Analytical solution from linear stability analysis in the limit *D* = 0 reveals three different regimes as function of the autocatalytic feedback strengths *μ* and *ζ*. **H**. Memo of interaction scheme highlighting parameters *α, μ, ζ* of Eqs. (1)-(2). Parameters: **A-C:** *λ* = 5000*/L*_sys_. For detailed list of model parameters, see Table S1 in SM.

To probe whether periodic patterns also form for a finite number of interacting molecules, we performed agent-based simulations with mean-field interactions. We introduce the total number of myosin filaments in the system as *λL*_sys_. Changing the number-density parameter *λ* allows to tune small-number fluctuations. The formal limit *λ* → ∞ recovers the deterministic mean-field model Eqs. (1-2). For efficient simulations, we introduce spatial bins of size Δ*x* and update the number of myosin filaments and Z-disc proteins in each bin according to Poisson birth and death processes with rates given by the product of *λ*Δ*x* and the respective binding and unbinding rates in Eqs. (1,2), see Supplemental Material for details.

This model assumes effective mean-field interactions, in which each myosin filament and Z-disc protein interacts with all other molecules at the same location. This assumption is a valid approximation in a dense, three-dimensional acto-myosin bundle.

Figure 3A shows snapshots of agent-based simulation at different simulation times with *λ* = 5000*/L*_sys_. At the start of simulation (*t* = 0), myosin filaments and Z-disc proteins are randomly distributed. As the pattern evolves in time, stable periodic patterns form. The number of filaments *M* in a bin can be translated into a concentration by *m* = *M/*(*λ* Δ*x*), see Fig. 3B. Analogous to the analysis of experimental data, we compute the autocorrelation function of myosin concentration, see Fig. 3C. To quantify the regularity of simulated patterns, we define a quality factor *Q* = −*π/* ln(*A*) from the amplitude *A* of the first Fourier-peak in these ACFs [46], see Fig. 3D. The quality factor *Q* increases with time and eventually saturates at a value *Q*_max_, reflecting the emergence of a stable pattern. The mean quality factor at steady-state *Q*_max_ increases with increasing number-density parameter *λ*, reflecting the decreasing impact of small-number fluctuations. In the formal limit *λ* → ∞, *Q*_max_ is expected to diverge to infinity as patterns become perfectly periodic, see Fig. 3E.

We emphasize that the formation of periodic patterns in model I is driven by the non-local interaction between myosin and Z-disc proteins, and does not emerge from a diffusion-driven instability as in classical Turing models [47]. Fig. 3F shows a phase diagram of model I as function of the non-local interaction strength *α* and the effective diffusion coefficient *D*, corroborating the fact that non-local interactions must outcompete the deleterious effects of diffusion to allow for the formation of periodic patterns. This remains true even if diffusion constants *D*_*m*_ ≠ *D*_*z*_ were different. In the limit *D* → 0, we can derive analytical conditions for the feedback parameters *ζ* and *μ* to predict whether stable periodic patterns, or no patterns or unstable patterns form, see Fig. 3G.

In conclusion, model I based on non-local interactions is capable of driving the spontaneous formation of periodic sarcomeric patterns, with sarcomere length set by the interaction length-scale of the non-local interactions.

### Sarcomeric pattern formation by tension-responsive catch-bonds

In addition to molecular interactions, mechanical tension was shown to be essential for sarcomere formation [8]. Myotubes are set under tension after they attach to tendon cells. Live-imaging in *Drosophila* revealed that periodic sarcomeric patterns emerge simultaneously across the entire length of initially unstriated muscle fibers, suggesting that the global level of mechanical tension in these fibers coordinates sarcomere assembly (concomitantly increasing nematic order and density of the actin filaments) [1, 8]. In light of this global role of mechanical tension guiding sarcomere assembly, it is tempting to speculate that local, position-dependent tension within nascent sarcomeres may also stabilize molecular interactions by modulating rates of unbinding. In particular, binding of *α*-actinin to actin filaments has been recently shown to display catch-bond behavior at the cortex of HeLa cells [34], and in reconstituted actin networks [35].

To explore this hypothesis, we introduce a second model, in which myosin filaments bound to anti-parallel actin filaments exert molecular tension on cross linked Z-disc proteins, see Fig. 4A. This provides an alternative non-local interaction of myosin filaments acting (indirectly) on Z-disc proteins. Together with the molecular interactions sketched in Fig. 2E, this defines model II of tension-dependent pattern formation. We make the simplifying assumption that translocation of actin filaments within the cross-linked actin bundle of the nascent myofibril can be neglected.

**FIG. 4.**
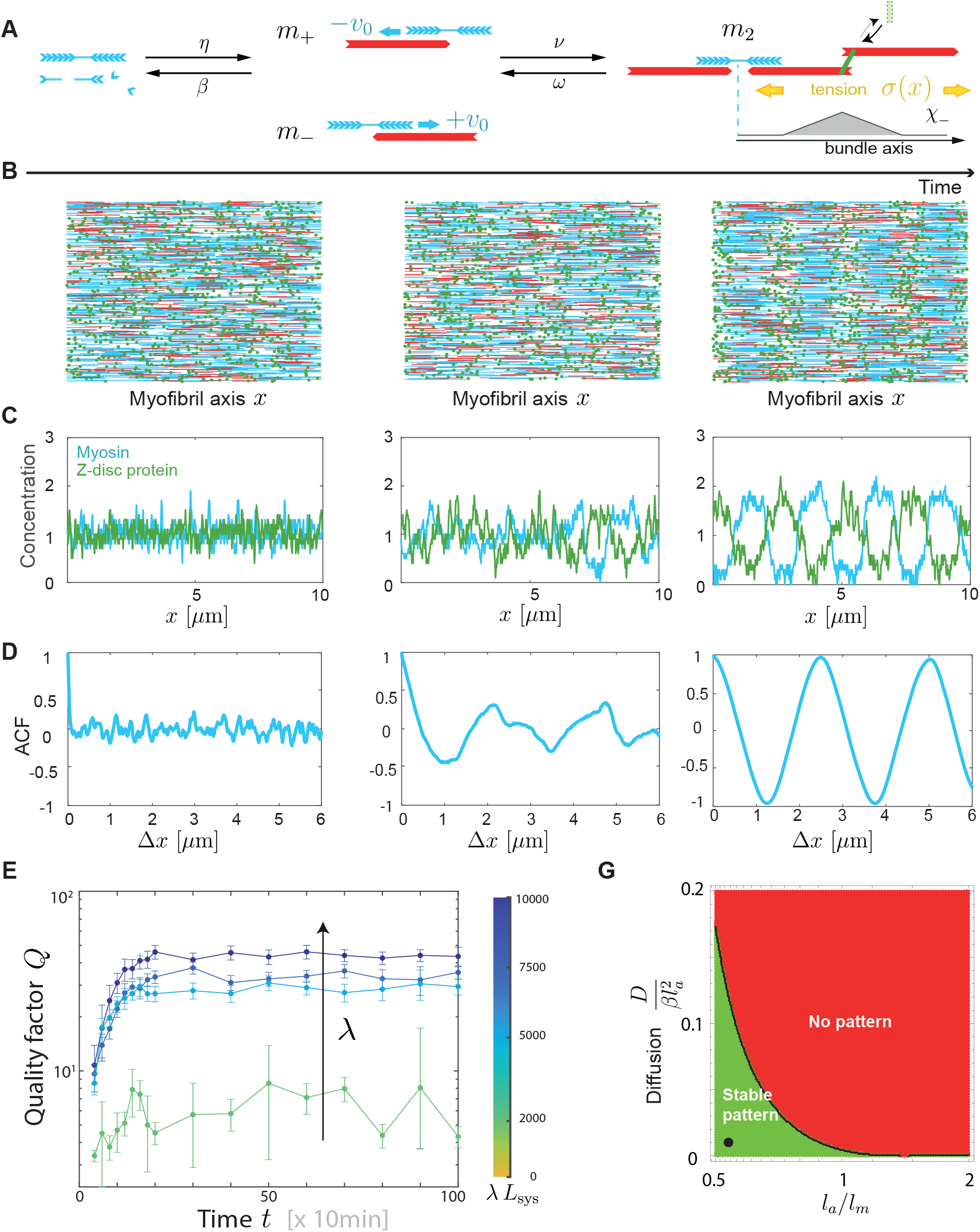
Pattern formation by tension-responsive catch-bonds. **A**. Tension-driven myofibrillogenesis: Bipolar myosin filaments can attach to polar actin filaments in different configurations labeled *m*_+_, *m*_−_, and *m*_2_ distinguished in Eqs. (5,6). Single-bound myosin with concentrations *m*_*±*_(*x*) are attached to actin filaments of only one polarity (referred to as + and −, respectively), and move towards the respective plus-end with velocity ∓*v*_0_. Double-bound myosin filaments with concentration *m*_2_(*x*) are attached simultaneously to actin filaments of opposite polarity, and thus do not move, but generate active tension *σ*(*x*). The Z-disc protein *α*-actinin is a catch-bond, i.e., its unbinding rate from actin decreases with increased tension. Together, this defines a second non-local interaction from myosin to Z-disc proteins. **B**. Snapshots of agent-based simulations of the mathematical model II at different simulation times (*t* = 0, 2, 100). Simulations of model II in Eqs. (5-7) were performed in one space dimension, and visualized in two dimensions for clarity with double-bound myosin (blue), Z-disc protein (green), polar actin filaments (red). **C**. Corresponding concentration profiles *m*_2_(*x*) and *z*(*x*) at time-points *t* = 0, 2, 100 for double-bound myosin (blue) and Z-disc protein (green). **D**. Corresponding ACFs for these concentration profiles. **E**. Quality factor *Q* characterizing the regularity of periodic myosin patterns as function of time *t* for different values of the number-density parameter *λ* scaling small-number fluctuations (mean*±*s.e.m., *n* = 10 simulation runs). **F**. Phase diagram showing regimes of stable patten formation as function of the relative length *l*_*a*_*/l*_*m*_ of actin filaments, and the effective diffusion coefficient of myosin filaments. Parameters: B-D: *λ* = 5000*/L*_sys_. For detailed list of model parameters, see Table S2 in SM.

In this model II, the structural polarity of actin filaments becomes important. In our one-dimensional model, there are two polarities of actin filaments, depending on whether their plus-end points to the left (+) or right (−). Correspondingly, we distinguish three different populations of myosin filaments bound to the scaffold of actin filaments, see Fig. 4A. Myosin filaments bound to actin filaments of only one polarity [with respective midpoint concentrations *m*_*±*_(*x*)] move to the respective plus-end of actin with velocity ∓*v*_0_. In contrast, myosin filaments bound to actin filaments of both polarities [with concentration *m*_2_(*x*)], do not move, but generate local tension *σ*(*x*) by pulling on actin filaments of opposite polarity. This local tension *σ*(*x*) can be expressed as a function of the concentration of double-bound myosin filaments *m*_2_(*x*) using interaction kernels *χ*_*±*_ that characterize the expected overlap of actin and myosin filaments, see also Fig. 4A

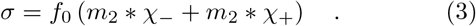

Specifically, in a mean-field description, the convolution kernels *χ*_*±*_ are normalized triangular pulse functions, with position and width set by the length of bipolar myosin filaments *l*_*m*_ and actin filaments *l*_*a*_, respectively.

To account for the tension-responsive catch-bond behavior of Z-disc proteins such as *α*-actinin, we assume that the unbinding rate *β*_*z*_ of Z-disc proteins decreases with increasing tension

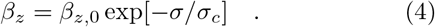

We can now formulate the minimal model II as a mean-field model that couples the concentrations of single-bound myosin *m*_*±*_(*x*), double-bound myosin *m*_2_(*x*), and Z-disc proteins *z*(*x*)

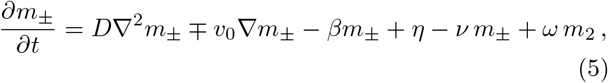

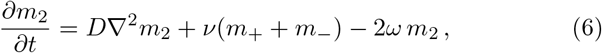

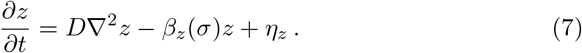

Eqs. (5-7) account for random motion modeled as effective diffusion with diffusion coefficient *D*, as well as unbinding and binding of myosin and Z-disc proteins to the scaffold of aligned actin filaments. Additionally, Eq. (5) contains a drift term that accounts for the active displacement of single-bound myosin along actin filaments. Single-bound myosin filaments that bind to a second actin filament of opposite polarity enter as a loss term −*νm*_*±*_ in Eq. (5) for *m*_*±*_(*x*), but as gain term in Eq. (6) for *m*_2_(*x*); analogously, *ωm*_2_ accounts for double-bound myosin filaments that unbind from actin filaments of one polarity, but stay bound to actin filaments of the opposite polarity, see Fig. 4A.

Analogous to model I, the binding and unbinding rate parameters *η* = *η*(*z*), *ν* = *ν*(*m*_2_), *ω* = *ω*(*m*_2_), *η*_*z*_ = *η*_*z*_(*z*) additionally depend on the local concentrations of myosin and Z-disc proteins, describing steric repulsion effects and autocatalytic binding, see Supplemental Material for details. The unbinding rate *β* for single-bound myosin contains an additional term 2*v*_0_*/L*_a_ that accounts for the fact that a single-bound myosin filament falls off the actin filament once it reaches its end.

Linear stability analysis of Eqs. (5-7) reveals again parameter regimes, for which the homogeneous steady state is unstable, and regular periodic patterns with alternating peaks of bound myosin and Z-disc proteins form.

Likewise, agent-based simulations, analogous to those for model I, reveal spontaneous sorting of myosin filaments and Z-disc proteins, see Fig. 4B, as well as Fig. 4C for concentration profiles at different simulation times. Amplitudes and wavelengths of emergent periodic patterns are comparable to those of model I. The wavelength of patterns is now set by the length of myosin and actin filaments.

From the auto-correlation functions of concentration profiles shown in Fig. 4D, we obtain again a quality factor that characterizes the regularity of patterns, see Fig. 4E. As expected, the quality factor increases with increasing number-density parameter *λ*, yet, as an important difference to model I, we find that no periodic patterns emerge if *λ* is too low with *λ* ≲ 1000*/L*_sys_.

Similar to small-number fluctuations, stochastic motion of molecules characterized by the effective diffusion coefficient *D* suppresses patterns, see Fig. 4F.

We emphasize that models I and II represent physically distinct mechanisms, yet share a similar feedback-logic. Correspondingly, they display a similar pattern forming behavior. Model II is slightly less robust than model I, reflected by smaller quality factors, and a collapse of periodic patterns already at intermediate values of the number-density parameter *λ*. Myofibrillogenesis *in vivo* likely uses a combination of models I and II.

## DISCUSSION

Here, we presented data on the early stages of myofibrillogenesis in *Drosophila melanogaster*, revealing a sequential ordering of the sarcomere components *α*-actinin, actin, muscle-specific myosin heavy chain, and the titin homologue Sallimus. Quantitative analysis using a new, tracking-free algorithm to compute auto- and cross-correlation functions shows that *α*-actinin, myosin and Sallimus establish periodic patterns with alternating localization first, while actin follows later.

Based on these observations, we propose two putative models of sarcomere self-assembly, which we formulate as minimal mathematical models. We propose that myosin and Z-disc proteins bind and unbind to a scaffold of parallelly aligned, but not yet polarity-sorted actin filaments, establishing periodic patterns as a consequence of auto-catalytic attachment, mutual interactions, and a negative feedback loop reflecting steric repulsion.

Our model I includes non-local interactions mediated by extended myosin filaments and the giant protein titin/Sallimus, which binds to bipolar myosin filaments at its C-terminus and is supposed to recruit Z-disc proteins such as *α*-actinin at its N-terminus. We thus assume in model I that Sallimus is incorporated already in its extended configuration, as has been shown to be the case in mature sarcomeres [5]. Sallimus would become mechanically strained later, possibly as consequence of an increase in sarcomere length. Intriguingly, previous experiments with genetically engineered titin demonstrated that varying titin length directly affects sarcomere length [28–30].

A modified model I, in which the roles of myosin and Z-disc proteins are flipped, where Z-disc proteins exert a non-local interaction on myosin, e.g., by recruiting Sallimus, which then stabilizes bound myosin, while myosin disfavors binding of Z-disc proteins due to steric hindrance, would yield analogous patterns by symmetry.

In a second model II, a similar non-local interaction results from the catch-bond behavior of Z-disc proteins such as *α*-actinin [34, 35] in response to local tension generated by myosin motor activity. Analogously, Z-disc protein complexes (e.g., Zasp52/*α*-actinin) or myosin itself could act as tension sensors [16].

Using agent-based simulations, we demonstrate the robustness of both mathematical models to small-number fluctuations, with a break-down of pattern formation below a critical number-density of about 50 myosin filaments per sarcomere unit in model I, but about 500 myosin filaments in model II. *In vivo* myofibrillogenesis could exploit a combination of model I and II, which may explain the high robustness of this process *in vivo*.

Our proposed feedback scheme does not represent a classical Turing mechanism with diffusion-driven instability [47]; instead, patterns emerge from non-local interactions, formally similar to previous work by Kondo et al. on zebrafish stripe patterns [48].

Based on data, our models assume that periodic actin patterns will become established later. A simple mechanism for subsequent actin ordering would be continuous actin turn-over, whereby new actin filaments become preferentially nucleated at nascent Z-discs (see also Fig. S8 in SM). Additionally, ectopic actin filaments that are not sufficiently crosslinked by Z-disc proteins may be moved by myosin motor activity and eventually either become captured at a Z-disc, or depolymerize. In particular, motion of these actin filaments may cause actin buckling, which could trigger accelerated depolymerization [49–51], see also previous studies that observe accelerated actin turnover as result of myosin activity [52–54]. Thus, preferential nucleation of new actin filaments at periodically positioned Z-disc precursors would ensure correct actin polarity within emerging sarcomeres, while the original scaffold of actin filaments with random polarity would be steadily remodeled and partially disassembled.

While it would be naïve to assume that myofibrillogenesis proceeds identically in all animals, we conceive that the physical principles proposed here could be conserved. For example, Sanger proposed a premyofibril model of myofibrillogenesis in which non-muscle myosin establishes periodic patterns first that later become replaced by muscle-specific myosin [55]. It is conceivable that our model may apply to a patterning of non-muscle myosin and Z-disc proteins. To validate our model, future cryo-electron tomography could assess the length and structural polarity of actin and myosin filaments, as recently done in mature sarcomeres [56, 57].

Our minimal mathematical models comprise effective parameters that coarse-grain biophysical parameters. The predicted patterning is robust and largely independent of specific parameters choices. Nonetheless, future fine-grained models should be quantitative and employ measured parameters, of which only some are known to date. Sarcomere assembly is a fast process in arthropods, spanning just 26 − 32 h APF for the *Drosophila* flight muscle.

In unstriated stress fibers, the length of actin filaments ranges from 0.5 − 2 *μ*m [58]; the distribution of actin filament length could be similarly disperse in early myofibrils. The length of bipolar myosin filaments in vertebrates equals *l*_*m*_ = 1.6 *μ*m in mature sarcomeres [59]. The density of myosin filaments in the cross-section of developing myofibrils at 32 h and 48 h APF estimated from electron micrographs equals approximately 150 *μ*m^−2^ [37] corresponding to about 35 myosin filaments per myofibrillar cross-section of 0.5 *μ*m diameter [39], or a value *λ* ∼ 150*/L*_sys_ of the number-density parameter in our model. During myofibrillogenesis, myosin expression is upregulated [12], and the number of individual myosin heavy chains per bipolar myosin filament increases with time [38, 39], which would correspond to a dynamic increase of an effective value of the number-density parameter *λ* in our model. Striated stress fibers with as low as 10-30 non-muscle myosin filaments in each myosin band were observed in cultured fibroblasts [60]. The stoichiometry between actin and myosin filaments was inferred as 3 : 1 in mature insect flight muscle [61, 62]. A sliding speed of non-muscle myosin of 0.15 *μ*m*/*min was observed in nascent striated stress-fibers [42]; we expect that the sliding speed *v*_0_ of muscle myosin during myofibril assembly stages is closer to this value than to the maximal speed 6 *μ*m*/*s of myosin filaments measured in mature fast skeletal muscle [63]. In mature myofibrils, each bipolar myosin filament (thick filament) contains 600 myosin motor heads [64], each of which can generate a maximal force of approximately 1 pN, though at low duty ratio *<* 0.1 [63, 65–67]. Each bipolar myosin filament is thus estimated to generate tensile forces in the range 1 − 10 pN [68].

Experiments in the actin cortex of HeLa cells indicate a critical tension of the catch-bond *α*-actinin of *γ*_*c*_ ≈ 1 nN*/μ*m [34]. Assuming a cortical thickness of *h* = 200 nm with actin network meshsize *a* = 50 nm, this value would correspond to a molecular force of *a*^2^*γ*_*c*_*/h* ≈ 10 pN.

FRAP experiments in beating cardiomyocytes revealed two actin populations with fast (∼1 min, 25%) and slow (*>* 30 min, 75%) turn-over, respectively [52]. Likewise, FRAP experiments in cultured quail myotubes indicated distinct populations of actin filaments and various Z-disc proteins including *α*-actinin, with typical turnover times ∼ 10 min [69]. For comparison, FRAP experiments in stress fibers indicate turnover rates of ≈ 0.5 min^−1^ for actin, and ≈ 3 min^−1^ for *α*-actinin [70].

The observation of two distinct actin populations with different turnover times in FRAP experiments of nascent myofibrils [52, 70] may reflect the initial scaffold of weakly crosslinked actin filaments and a subsequent population of anchored actin filaments that may have been polymerized *de novo* at nascent Z-discs as proposed here.

Generally, the exchange dynamics of sarcomeric components seems to slow down during myofibril maturation [69]. Turnover of titin/Sallimus in mature *Drosophila* larval muscle was slow with turn-over times *>* 30 min [27], consistent with previous results for titin turnover in cultured mouse myocytes of several hours [71].

Assuming a value of 0.1 min^−1^ for binding and unbinding rates in our mathematical model would convert the time for the formation of periodic sarcomeric patterns from 20 simulation time units to 3 h, which is at least consistent with observations. Similarly, myosin speed *v*_0_ in model II would correspond to 0.2 *μ*m*/*min.

Our mathematical model makes testable predictions: Knock-out of myosin should result in the formation of Z-disc protein aggregates. In fact, such Z-bodies may have already been observed [15, 45, 72]. Knock-out of essential Z-disc proteins or over-regulation of autocatalytic attachment of myosin should likewise result in the formation of myosin stacks. We thus consider it likely that Z-bodies or myosin stacks observed in previous studies with atypical myogenic cells, mutants, or harsh pharmacological treatments [15, 16, 42, 43, 45] may not represent “building blocks” of sarcomere self-assembly in the literal sense that such building blocks are physically stitched together to form myofibrils, but instead “logical building blocks” that manifest key feedback mechanisms underlying sarcomere assembly.

## Supporting information

Supplemental text including 8 supplemental figures

## Acknowledgment

FS and BMF acknowledge financial support from the Human Frontiers Science Program (HFSP, grant: RGP0052); FS acknowledges support from the Centre National de la Recherche Scientifique (CNRS), the European Research Council under the European Union’s Horizon 2020 Programme (ERC-2019-SyG 856118), the excellence initiative Aix-Marseille University A*MIDEX (ANR-11-IDEX-0001-02), the France-BioImaging national research infrastructure (ANR-10-INBS-04-01) and by funding from France 2030, the French Government program managed by the French National Research Agency (ANR-16-CONV-0001) and from Excellence Initiative of Aix-Marseille University - A*MIDEX (Turing Centre for Living Systems).

BMF acknowledges financial support from the Deutsche Forschungsgemeinschaft (DFG, German Research Foundation) under Germany’s Excellence Strategy - EXC-2068-390729961, as well as through a Heisenberg grant of the DFG (FR3429/4-1,4-2).

