## Supplemental text including 8 supplemental figures for "Theory of sarcomere assembly inferred from sequential ordering of myofibril components"

### Supplemental Material

*Experimental methods.* *Drosophila Mhc*[*weeP26*](*Mhc*-GFP) larvae were grown at 27°C and collected at the time of puparium formation (white colored pupae). Pupae were then maintained at 27°C for at least 22 hours and up to 32 hours After Puparium Formation (APF), which corresponds to the time window of sarcomere assembly in *Drosophila* Indirect Flight Muscles at that temperature [8, 38]. At each time point (22, 24, 26, 28, 30 and 32 hours APF), pupae were fixed with 4% paraformaldehyde in PBS-T (PBS + 0.3% Triton X-100) for 30 min at room temperature. Pupae were then washed twice in PBS-T for at least 10 min, and, still in PBS-T, thoraxes were dissected and cut in half as described in [8] to visualize Indirect Flight Muscles. Staining was done in PBS-T: actin was labeled with Phalloidin Alexa-568 (Thermo Fisher, 1:500), titin/Sallimus and $\alpha$-actinin were labeled with the following fluorescently labeled nanobodies [27] Sls-Nano2-DyLight405 and Actn-Nano62-Atto647, respectively. Samples were stained overnight at 4°C, then washed 3 times (3x10 min) in PBS-T and mounted in SlowFadeTM Gold Antifade (Thermo Fisher). Samples were imaged with a Zeiss LSM880 confocal microscope using an 100× objective in the AiryScan mode (R-S), with a 2.5× zoom. We processed the Airyscan raw data within the software ZEN using the default (automatically calculated) strength value.

The experiment was performed twice and images were acquired from in total 7 different animals. Staining for $\alpha$-actinin had been added in the second experimental run, which corresponds to 4 different animals.

*Model parameters.* For all simulations, non-dimensionalized parameters were used. Tables S1, S2, S3 list the model parameters used for Fig. 3, Fig. 4, as well as numerical parameters, respectively.

| Model I | | | |
|---|---|---|---|
| Description | Parameter | Value | Unit |
| Steady-state concentrations | $m^*$, $z^*$ | 1 | - |
| Interaction length | $l_m/2 + l_s$ | 1 | $\mu$m |
| Effective diffusion coefficients | $D$, $D_m$, $D_z$ | 0.01 | $0.1\,\mu\text{m}^2/\text{min}$ |
| Unbinding rate | $\beta$ | 1 | $0.1\,\text{min}^{-1}$ |
| Feedback strengths | $\zeta$, $\mu$ | 0.85 | - |
| Non-local interaction strength | $\alpha$ | 1 | - |
| Number-density parameter | $\lambda$ | $20 - 10^3$ | $\mu\text{m}^{-1}$ |

TABLE S1. Model parameters used for the simulations of Eqs. (1-2) for model I shown in Fig. 3.

| Model II | | | |
|---|---|---|---|
| Description | Parameter | Value | Unit |
| Steady-state concentrations | $m_{\pm}^{*}$, $m_{2}^{*}$, $z^{*}$ | 1 | - |
| Steady-state tension | $\sigma^{*}$ | 2 | $10\,\mathrm{pN}/\mu\mathrm{m}$ |
| Myosin length | $l_m$ | 1.6 | $\mu$m |
| Actin length | $l_a$ | 1 | $\mu$m |
| Effective diffusion coefficients | $D_{\pm}$ ,$D_2$, $D_z$ | 0.01 | $0.1\,\mu\mathrm{m}^2/\mathrm{min}$ |
| Myosin speed | $v_0$ | 2 | $0.1\,\mu\mathrm{m}/\mathrm{min}$ |
| Binding time scales | $\tau = \tau_z$ | 0.15 | 10 min |
| | $\tau_2$ | 1 | 10 min |
| Feedback exponents | $\epsilon_z$ | 0.5 | - |
| | $\epsilon_\alpha$ | 1 | - |
| | $\epsilon$ | 1 | - |
| Critical tension | $\sigma_c$ | 1 | $10\,\mathrm{pN}/\mu\mathrm{m}$ |
| Myosin force | $f_0$ | 1 | 10 pN |
| Number-density parameters | $\lambda$, $\lambda_a$ | $20 - 10^3$ | $\mu\mathrm{m}^{-1}$ |

TABLE S2. Model parameters used for the simulations of Eqs. (5-7) for model II shown in Fig. 4.

| Numerical parameters | | | |
|---|---|---|---|
| Description | Parameter | Value | Unit |
| System size | $L_{\mathrm{sys}}$ | 10 | $\mu$m |
| Spatial bin size | $\Delta x$ | 0.02 | $\mu$m |
| Time step | $\Delta t$ | 0.01 | 10 min |
| Regularization length for spatial coarse-graining | $\varepsilon$ | 0.08 | $\mu$m |

TABLE S3. Numerical parameters used for agent-based simulations of model I and II shown in Figs. 3, 4.

*Details on tracking-free algorithm for correlation functions.* Fig. S1 shows a more detailed description of the tracking-free algorithm used to compute auto- and cross-correlation functions from micrographs of early myofibrillogenesis. In a first step, pre-processing was applied to the raw images to enhance relevant features, by adaptive filtering by using a Gauss-filter using the Matlab function `imgaussfilt` (setting $\sigma = 2$), followed by a top-hat transformation. using the function `imtophat` (setting $\sigma = 28$). Next, images were binarized using a fixed threshold manually set for each image.

To compute local nematic order and local linescans, moving regions-of-interest (ROI) of size $30 \times 30$ pixel were used. Only ROIs with a minimum of 25% of white pixels were included. ROIs close to the image boundaries were discarded to ensure that local linescans (which are larger than the ROIs) do not exceed the boundaries of the image.

To determine the local nematic direction in each ROI, which serves as proxy for the common local tangent of myofibrils in each ROI, a steerable filter (using free software from Francois Aguet[1]) was applied to the Sallimus channel of each image. We decided to use the Sallimus channel to determine the local nematic direction, as this channel turned out to provide the clearest nematic signals, whereas, e.g., the actin channel gave a more diffuse signal at $22-26$ h APF. The nematic directions were determined by monitoring the maximum response of this steerable filter ($0-180°$), implemented as convolution with the directed second derivative of a Gaussian kernel, which is particularly sensitive to edges and fiber-like structures. For each ROI, linescans centered at the center of the ROI of total length $5\,\mu$m and width 1 pixel parallel to the local nematic direction of the ROI were computed. Note that the linescans are longer than the ROI size.

From the intensity profiles obtained by these linescans, individual, unbiased auto-correlation functions (ACF) and cross-correlation functions (CCF) were computed and normalized by the respective variances. These normalized correlation functions were then averaged over all ROIs; see Fig. 1C for a typical example.

---

[1] http://www.francoisaguet.net/software.html, inspired by M. Jacob and M. Unser, "Design of Steerable Filters for Feature Detection Using Canny-Like Criteria", IEEE Trans. Pattern Anal. Mach. Intell., 26(8), pp. 1007-1019, 2004

Note that in several images an intensity gradient was present, which can cause auto-correlation functions to drop below zero, as can be seen, e.g., in Fig. 1C for the case of 22 h.

A minimal version of the tracking-free algorithm is available at

`https://github.com/BDehapiot/BDProject_FDomMuscle_2D`.

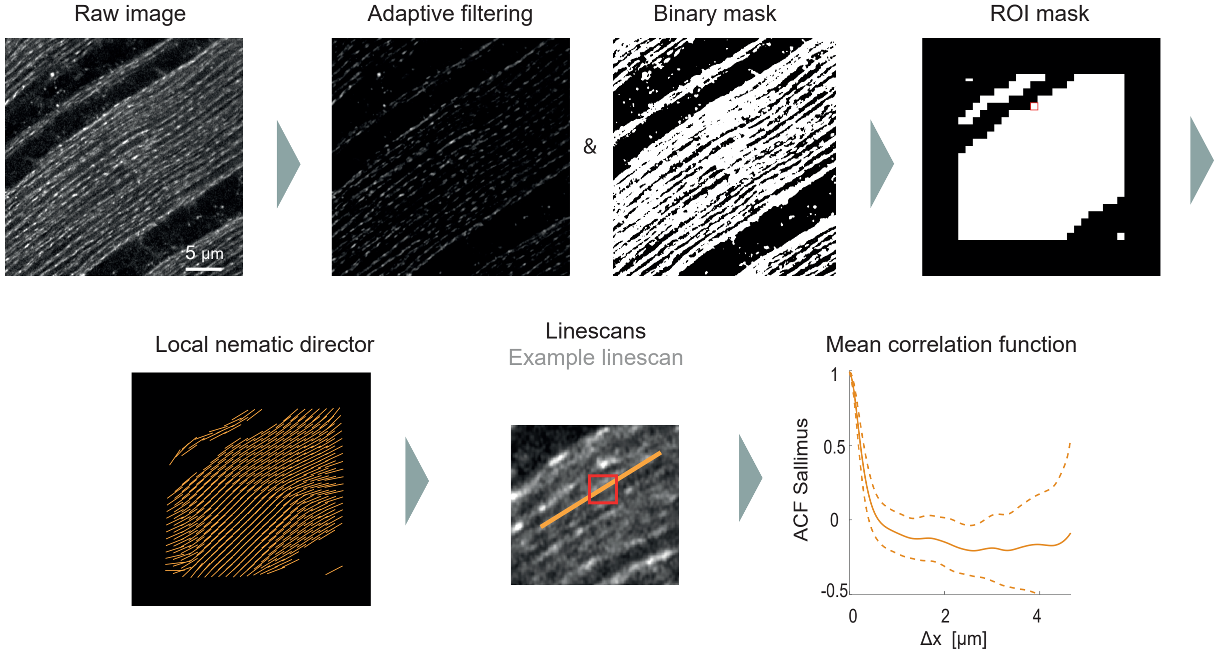


FIG. S1. **Tracking-free algorithm for correlation functions.** *Step 1:* Pre-processing of raw images by adaptive filtering and binarization. Mask of selected ROIs (example ROIs shown as red square). *Step 2:* Local nematic director determined for each ROI using a steerable filter. *Step 3:* Linescans are computed for each ROI along this nematic director. *Step 4:* Finally, normalized correlation functions are computed from these linescans and averaged over all ROIs (mean±s.d.).

*Additional correlation functions.* To complement Fig. 1C, Fig. S2 shows all correlation functions for all measured time-points.

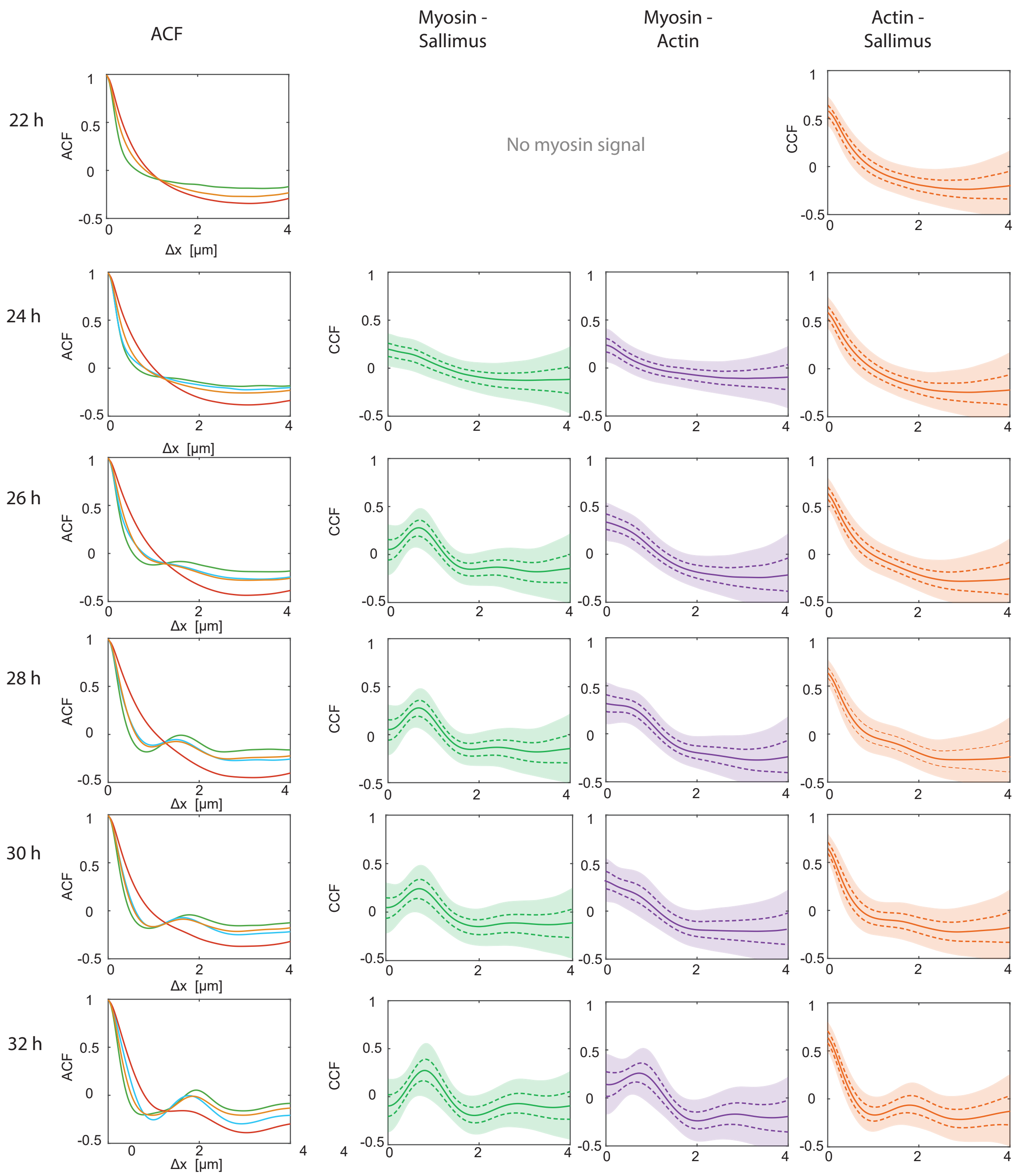


FIG. S2. Autocorrelation and cross-correlation functions analogous to Fig. 1 for all combinations of channels for myosin, actin, Sallimus, for each measured time-point of early myofibrillogenesis.

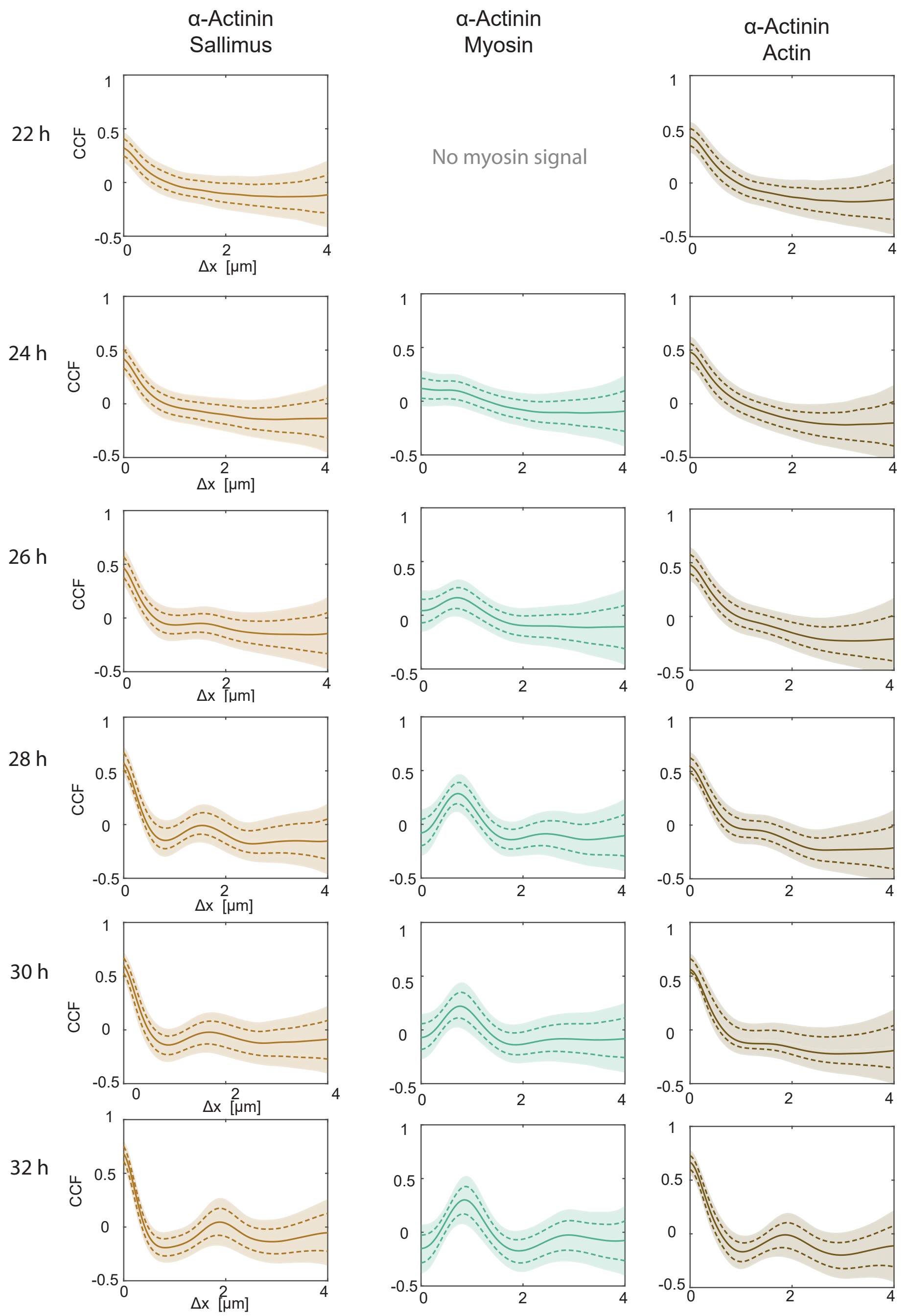


FIG. S3. Cross-correlation functions analogous to Fig. 1 for all combinations of channels with $\alpha$-actinin, for each measured time-point of early myofibrillogenesis.

To systematically investigate micrographs displaying advanced myofibrillogenesis, we automatically filtered the top 5% of all ROIs from every z-stack, for which the Fourier peak in the ACF of Sallimus was maximal (determined from fitting a `sin`-function to the ACF around the expected peak position). Fig. S4 shows ACFs for actin, myosin, and Sallimus averaged over these selected ROIs. As expected, Fourier peaks become more pronounced for all channels, as compared to the mean over all data as shown in Fig. S2. In particular, the ACF for actin exhibits a Fourier peak already at 24 h in Fig. S4, but only at 26 h in Fig. S2. For Sallimus, the Fourier peak is much more pronounced at 26 h, and even a second Fourier peak becomes visible at 28 h. However, the relative timing of sarcomeric pattern formation is identical, irrespective of whether the most advanced ROIs are considered as in Fig. S4, or an average is taken over all ROIs as in Fig. S2, with a time-shift of at most 2 h between both.

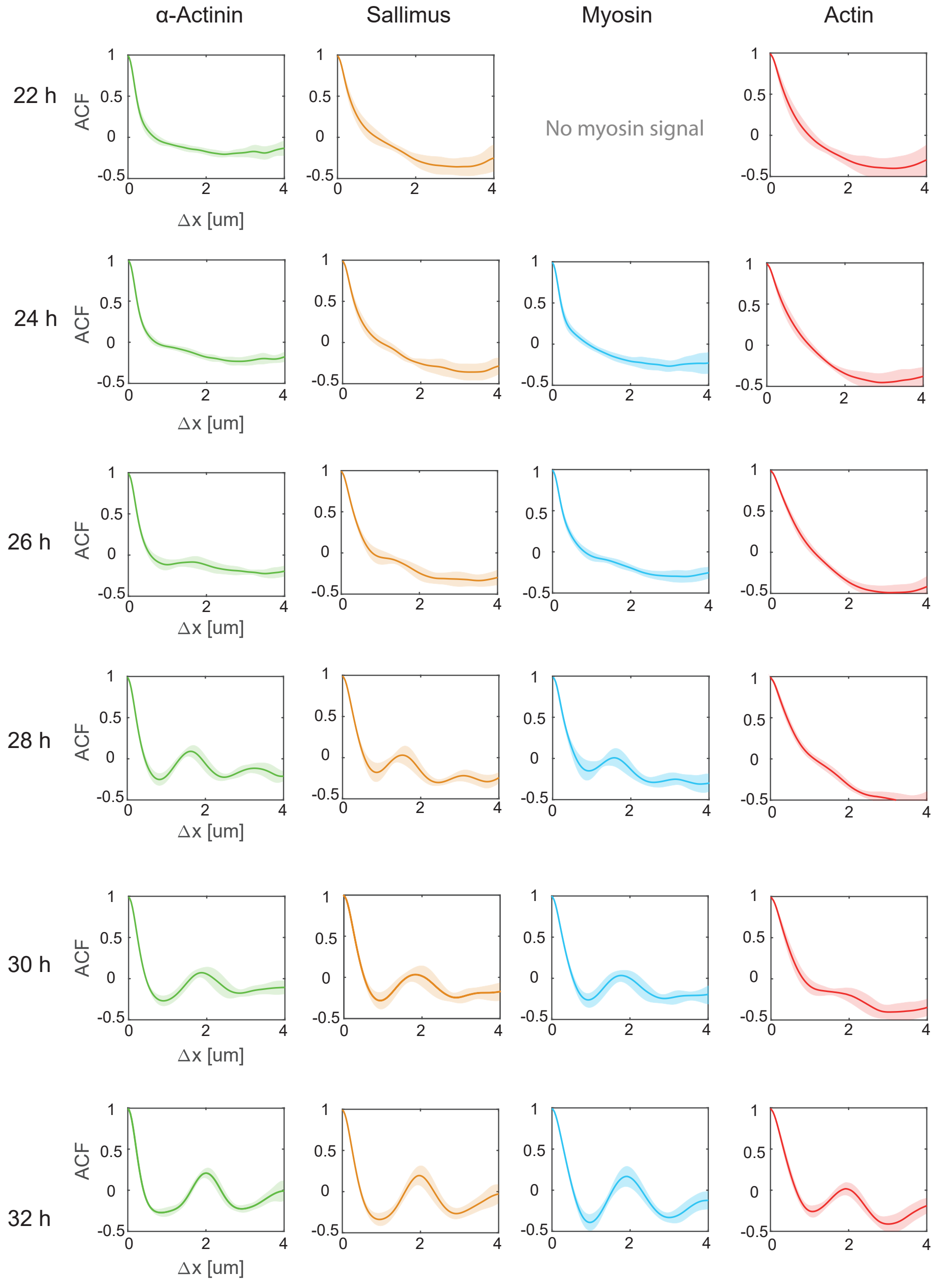


FIG. S4. Normalized auto-correlation functions for actin, myosin, and Sallimus (±s.d.), averaged over selected ROIs corresponding to the top 5% of most advanced ROIs.

*Sarcomere size at early stages of myofibrillogenesis.* The position $\Delta x^*$ of the principal Fourier peak of ACFs of noisy periodic patterns indicates the pattern length scale. For noisy patterns, the amplitude $A$ of this Fourier peak is a measure for the regularity of patterns, which we report in terms of the quality factor $Q = -\pi/\ln(A)$, see main text. Reduced amplitudes $A$ and correspondingly reduced quality factors $Q$ result in a small but systematic shift of the position $\Delta x^*$ of the principal Fourier peak relative to the true pattern length scale $L$. Typical ACFs can be approximated as a damped oscillation, where $Q$ sets the damping

$$\mathrm{ACF}(\Delta x) \approx \cos\left(\frac{2\pi\Delta x}{L}\right)\exp\left(-\frac{2\pi\,\Delta x}{LQ}\right)\ . \tag{S1}$$

(Indeed, Eq. (S1) becomes exact for a minimal model of noisy periodic patterns given by $I(x) \sim \cos(\varphi(x))$ where the phase variable $\varphi(x)$ of periodic patterns obeys the stochastic differential equation $d\varphi(x)/dx = 2\pi/L + \xi(x)$, where $\xi(x)$ denotes Gaussian white noise with $\langle \xi(x)\xi(x')\rangle = 2\pi/(QL)\,\delta(x-x')$.)

The peak position $\Delta x^*$ of the ACF given in Eq. (S1) is related to the pattern length scale $L$ by

$$L = \Delta x^* \left(1 + \frac{1}{2\pi}\tan^{-1}\left(\frac{\ln(A)}{\pi}\right)\right)\ . \tag{S2}$$

Fig. S5 reports sarcomere length $L$ inferred from the ACFs of all four sarcomeric proteins using both the correction formula Eq. (S2) as well as the uncorrected Fourier-peak position $\Delta x^*$. Eq. (S2) gives more consistent results compared to the uncorrected Fourier-peak position $\Delta x^*$.

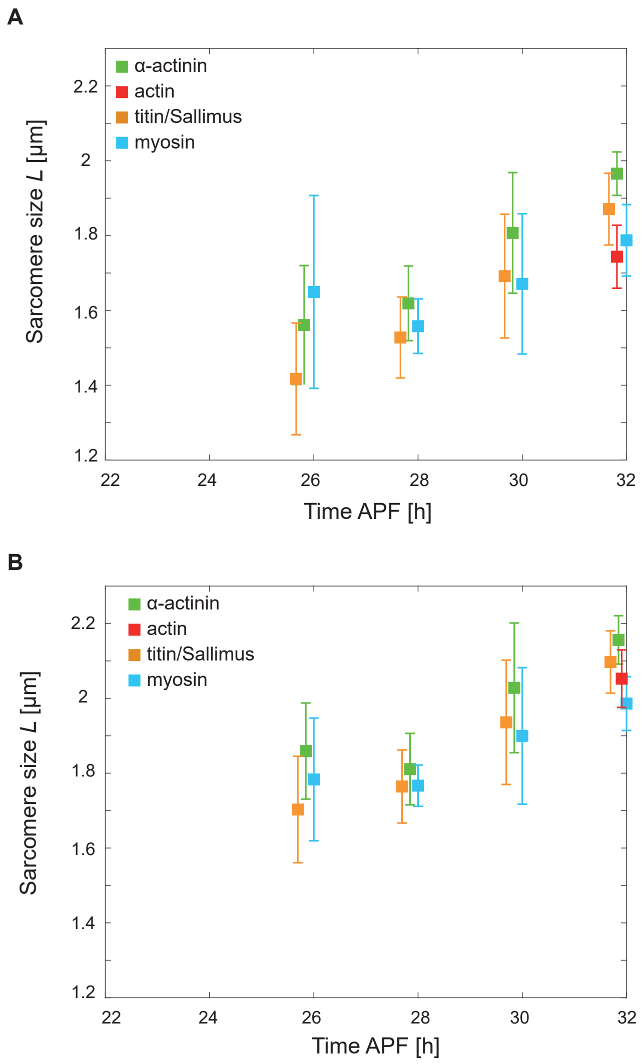


FIG. S5. **A.** Uncorrected sarcomere size, measured from the position $\Delta x^*$ of the first Fourier-peak of the ACFs for all four sarcomeric proteins. **B.** Corrected sarcomere size $L$ for all four channels using Eq. (S2).

*Agent-based simulations.* To probe the robustness of the proposed models with respect to small-number fluctuations, we performed agent-based simulations with a mean-field approximation for interactions. For efficient simulations of model I, we introduced spatial bins of size $\Delta x$ and update the number $M_i$ and $Z_i$ of myosin filaments and Z-disc proteins in each bin with center $x_i = i\Delta x$ according to Poisson birth and death processes with rates given by the product of $\lambda\Delta x$ and the binding and unbinding rates in Eqs. (1) and (2). Note that the bin counts $M_i$ track the midpoints of extended myosin filaments. To evaluate the mathematical expressions for these rates, the concentrations $m(x)$ and $z(x)$ in these expressions are replaced by the stochastic number densities $M_i * \chi_\varepsilon/(\lambda\Delta x)$ and $Z_i * \chi_\varepsilon/(\lambda\Delta x)$, respectively, where $\chi_\varepsilon$ is a regularization kernel chosen as a Gaussian kernel with variance $\varepsilon^2$ that ensures that simulation results are independent of the choice of bin size $\Delta x$.

To simulate time-dynamics, we employ an Euler scheme with fixed time-step $\Delta t$. In each time step, we update the number $M_i$ and $Z_i$ of myosin filaments and $Z$-disc proteins in each bin, respectively, by adding and subtracting Poisson distributed random variables with mean (and variance) given by the expected number of binding and unbinding events, as well as the expected number current of diffusing molecules between neighboring bins.

Specifically, the update step for the number $M_i$ of myosin filaments in bin $i$ analogous to Eq. (1) reads

$$M_i(t_{j+1}) = M_i(t_j) + J_{M,i-1\to i} + J_{M,i+1\to i} - J_{M,i\to i-1} - J_{M,i\to i+1} - \Delta M_{\mathrm{unbind},i} + \Delta M_{\mathrm{bind},i} \quad . \tag{S3}$$

To determine the numbers $\Delta M_{\mathrm{unbind},i}$ of myosin filaments that will unbind from each bin in the next time-step, we first determine the total number of myosin filaments $\Delta M_{\mathrm{unbind}}$ that will unbind in the next time-step in the entire system as a Poisson distributed random number with mean $\beta\, dt \sum_i M_i$, and then select the respective number of integer indices marking the filaments to be removed from each bin. The unbinding of Z-disc proteins was calculated analogously, with total number $\Delta Z_{\mathrm{unbind}}$ of Z-disc proteins to be removed in the next time-step drawn as Poisson random variable with mean $\beta\, dt \sum_i Z_i$. We ensure that never more molecules than are present in the system can be removed. Accordingly, bin counts stay always non-negative.

The number $\Delta M_{\mathrm{bind},i}$ of myosin filaments that bind in bin $i$ in the next time-step is likewise drawn as a Poisson random variable

$$\Delta M_{\mathrm{bind},i} \sim \mathcal{P}\left(\beta\, m^* \exp\left[\mu\left(m_i - z_i - (m_i - m^*)^2\right)\right]\right) \quad , \tag{S4}$$

where the effective local concentrations $m_i$ and $z_i$ are determined from the bin counts as follows

$$m_i = (\lambda\Delta x)^{-1} \sum_k M_{i-k}\chi_{\varepsilon,k} \text{ and } z_i = (\lambda\Delta x)^{-1} \sum_k Z_{i-k}\chi_{\varepsilon,k} \quad . \tag{S5}$$

The regularization kernel $\chi_{\varepsilon,k}$ was chosen as a Gaussian kernel with mean 0 and variance $\varepsilon^2 = 0.08$, normalized to 1.

The deterministic diffusion current $J_{M,i-1\to i}$ of myosin filaments from bin $i-1$ to bin $i$ is given by

$$J_{M,i-1\to i} = M_{i-1} \int_{\Delta x/2}^{\infty} dx\, \frac{1}{\sqrt{4\pi\, D\Delta t}} \exp\left[\frac{-x^2}{4D\Delta t}\right] \quad . \tag{S6}$$

The integral on the right-hand side is closely related to the complementary error function $\mathtt{erfc}(x(4D\,\Delta t)^{-1/2})$. The other diffusion currents in Eq. (2) are defined analogously. We tested in preliminary simulations that accounting for diffusive currents to next-to-nearest-neighbor bins does not change results.

The update step for the number $Z_i$ of Z-disc proteins in bin $i$ corresponding to Eq. (2) reads

$$Z_i(t_{j+1}) = Z_i(t_j) + J_{Z,i-1\to i} + J_{Z,i+1\to i} - J_{Z,i\to i-1} - J_{Z,i\to i+1} - \Delta Z_{\mathrm{unbind},i} + \Delta Z_{\mathrm{bind},i} \quad . \tag{S7}$$

While unbinding rate and diffusion currents were calculated analogously to those for myosin, the number $\Delta Z_{\mathrm{bind},i}$ of Z-disc proteins that bind to bin $i$ in the next time-step was drawn from a Poisson distribution as

$$\Delta Z_{\mathrm{bind},i} \sim \mathcal{P}\left(\beta\, z^* \exp\left[\zeta\left(\alpha\,(m_{\chi,i} - m^*) + (z_i - z^*) - (z_i - z^*)^2\right)\right]\right) \tag{S8}$$

with $z_i$ as described in equation (S5), and $m_{\chi,i} = m_i * \chi_{\mathrm{m}} = (\lambda\Delta x)^{-1} M_i * \chi_{\mathrm{m}}$.

The same numerical methods were used for the agent-based simulations of model II shown in Fig. 4. The mathematical model II additionally explicitly accounts for actin filaments of specific position and polarity. For this, the number $A_i^{\pm}$ of actin filaments in each bin $i$ was drawn from a Poisson random distribution with mean $A^* = \lambda_a dx\, a^*$, for each actin polarity. Note that these bin counts again track the midpoints of extended actin filaments of length $l_a$. To account for local fluctuations in actin density, the rates of myosin filaments and Z-disc proteins binding to the actin bundle were multiplied with a factor $A * \chi_a / A^*$, where $\chi_a = \theta(l_a/2 - x)\theta(x + l_a/2)/l_a$ is a normalized, symmetric pulse-function of width $l_a$ that accounts for the non-zero length of actin filaments.

*Extended model I.* A slightly more general version of model I given in Eqs. (1) and (2) can be given as follows

$$\frac{\partial m}{\partial t} = D_m \nabla^2 m - \beta_m m + \beta_m m^* \exp\left[\mu\left(m - z - \gamma_m (m - m^*)^2\right)\right] \tag{S9}$$

$$\frac{\partial z}{\partial t} = D_z \nabla^2 z \quad - \beta_z z \quad + \beta_z z^* \exp\left[\zeta\left(\alpha\,(m_\chi - m^*) + (z - z^*) - \gamma_z (z - z^*)^2\right)\right] \quad , \tag{S10}$$

where we now allow for different diffusion coefficients $D_m$ and $D_z$, as well as different unbinding rates $\beta_m$ and $\beta_z$, and saturation parameters $\gamma_m$ and $\gamma_z$.

Linearizing this extended model I given by Eqs. (S9,S10) around its spatially homogeneous steady state $m(x) \equiv m^*$, $z(x) \equiv z^*$ gives the following Jacobian matrix as function of wavevector $k$

$$J(k) = \begin{pmatrix} \beta_m(\mu - 1) - D_\mathrm{m} k^2 & -\beta_m \mu \\ \beta_z \zeta \alpha\, \mathcal{F}(\chi_m) & \beta_z(\zeta - 1) - D_z k^2 \end{pmatrix} \quad . \tag{S11}$$

Here, $\mathcal{F}(\chi_m)$ denotes the Fourier transformation of the non-local interaction kernel $\chi_m$.

A linear stability analysis reveals that the extended model I can exhibit spontaneous pattern formation as a result of the non-local interaction, but does not exhibit any diffusion-driven instability for any general choice of parameters. (As a technical point, it is possible to realize a diffusion-driven instability with this extended model, but only for a very restricted parameter set.) Fig. S6B shows the dispersion relation for the parameter set used in Fig. 3.

The critical wavevector $k_c = 2\pi/L_c$ at which the largest real part of the eigenvalues of $J(k)$ becomes maximal is indicative of the wavelength of emergent patterns. Fig. S6C displays the critical wavelength $L_c$ as function of the length-scale of the non-local interaction, revealing a slope of approximately 2. This corroborates the observation that the size of sarcomere units in simulations accommodates two times the length-scale of non-local interactions. Note that for a finite system with system size $L_\mathrm{sys} = 10$ and periodic boundary condition as used in Fig. 3, we will only be able to observe an integer number of pattern units, in this case four or five, while the critical wavelength approximately equals $\approx 2.1$ for the parameters used.

We confirmed that choosing different diffusion coefficients $D_m$ and $D_z$, or unbinding rates $\beta_m$ and $\beta_z$ lead to a dynamics very similar to the one reported in Fig. 3.

The saturation parameters $\gamma_m$ and $\gamma_z$ do not influence the pattern-formation instability, but affect the shape of steady-state patterns. In particular, these parameters can be tuned such that steady-state patterns feature sharper Z-disc proteins bands reminiscent of nascent Z-discs, see Fig. S6A.

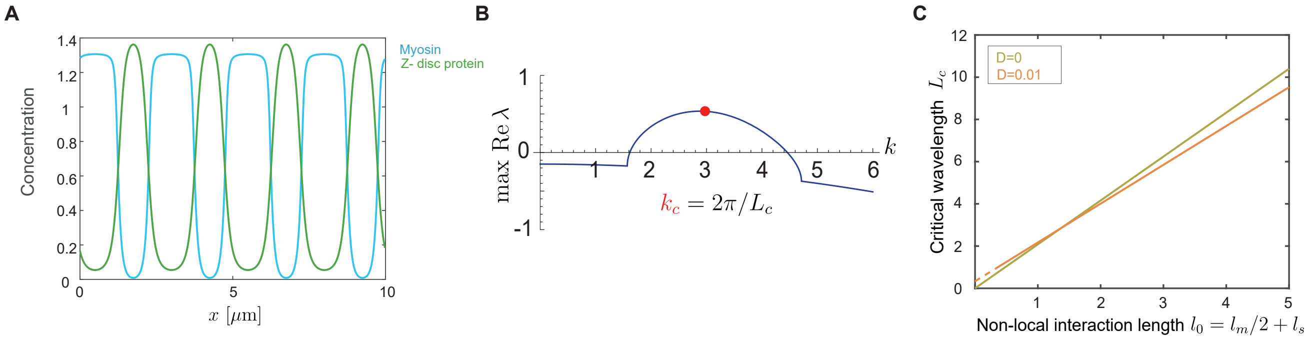


FIG. S6. **A.** Mean-field simulation of the extended model I given by Eqs. (S9-S10) analogous to Fig. 3, but with different saturation parameters for myosin and Z-disc protein, respectively, $\gamma_m = 10$ and $\gamma_z = 1.5$. **B.** Dispersion relation from linear stability analysis for model I. **C.** Critical wavelength $L_c$ as function of the length-scale $l_0$ of the non-local interaction between myosin and Z-disc proteins. Parameters as in Table S2, unless stated otherwise.

*Extended Model II.* A slightly more general version of model II given in Eqs. (5-7) can be given as follows

$$\frac{\partial m_\pm}{\partial t} = D_\pm \nabla^2 m_\pm \mp v_0 \nabla m_\pm - \beta m_\pm + \eta - \nu\, m_\pm \qquad + \omega\, m_2 \quad , \tag{S12}$$

$$\frac{\partial m_2}{\partial t} = D_m \nabla^2 m_2 \qquad + \nu(m_+ + m_-) - 2\omega\, m_2 \quad , \tag{S13}$$

$$\frac{\partial z}{\partial t} = D_z \nabla^2 z \qquad - \beta_z(\sigma) z + \eta_z \quad , \tag{S14}$$

where we use short-hand for concentration-dependent binding and unbinding rates

$$\begin{aligned}
\beta &= \beta_0 + \frac{2v_0}{l_a} \quad , \\
\eta &= \eta_0 \exp[-\epsilon_\eta z] \quad , \\
\nu &= \nu_0 \exp[\epsilon_\nu m_2 - \gamma_m (m_2 - m_2^*)^2] \quad , \\
\omega &= \beta_0 \exp[-\epsilon_\omega m_2] \quad , \\
\epsilon &= \epsilon_\nu + \epsilon_\omega \quad , \\
\eta_z &= \eta_{z,0} \exp[\epsilon_z z - \gamma_z (z - z^*)^2] \quad , \\
\beta_z &= \beta_{z,0} \exp[-\sigma/\sigma_c] \quad .
\end{aligned} \tag{S15}$$

The quadratic terms in the exponentials represent saturation effects and ensure finite amplitudes of concentration profiles at steady state. The saturation parameter $\gamma_m$ and $\gamma_z$ can be introduced similar to model I. Because we aim for symmetric patterns, we set $\gamma_m = \gamma_z = 1$.

The unbinding rate $\beta(\sigma)$ of Z-disc proteins depends on local mechanical tension $\sigma(x)$ as follows. Our minimal model for local mechanical tension given in Eq. (3) comprises interaction kernels $\chi_+$ and $\chi_-$ that characterize the overlap of myosin filaments with actin filaments. These interaction kernels are represented by normalized triangular pulse functions, depending on the lengths $l_m$ and $l_a$ of myosin and actin filaments, respectively

$$\chi_\pm(x) = \frac{1}{l_a^2} \begin{cases} l_a - \left| x \mp \frac{l_m}{2} \right| & \text{if } |x \mp l_m/2| < l_a \\ 0 & \text{else} \end{cases} \quad . \tag{S16}$$

The steady-states of Eqs. (5)-(7) for model II read

$$m_\pm^* = \frac{\eta^*}{\beta} = \eta^* \tau \tag{S17}$$

$$m_2^* = \frac{\gamma}{\omega} m_\pm^* \tag{S18}$$

$$z^* = \eta_z^* \frac{1}{\beta_z^*(\sigma^*)} \tag{S19}$$

$$\sigma^* = 2 f_0 m_2^* \quad . \tag{S20}$$

The concentration-dependent binding and unbinding rates at steady-state read

$$\eta^* = \beta^* m_\pm^* = m_\pm^* \frac{1}{\tau} \tag{S21}$$

$$\nu^* = \frac{1}{\tau_2} \frac{m_2^*}{m_\pm^*} \tag{S22}$$

$$\beta_z^* = \beta_{z,0} \exp(-\sigma^*/\sigma_c) \quad , \tag{S23}$$

where we introduced effective time-scales $\tau = 1/\beta$, $\tau_2 = 1/\omega^*$.

We performed a linear stability analysis around the spatially homogeneous steady state given by $m_\pm(x) \equiv m^*$, $m_2(x) \equiv m_2^*$, $z(x) \equiv z^*$, $\sigma(x) \equiv \sigma^*$. Note that the quadratic terms in the exponents in the definitions of the binding rates $\nu$ and $\eta_z$ do not affect this linearization. We thus obtain as an intermediate step towards linearization

$$\begin{aligned}
\frac{\partial m_\pm}{\partial t} =& \frac{1}{\tau} \left( m_\pm^* \exp[-\epsilon_\eta (z - z^*)] - m_\pm \right) + \frac{1}{\tau_2} m_2 \exp[-\epsilon_\omega (m_2 - m_2^*)] \\
&- \nu^* m_\pm \exp[\epsilon_\nu (m_2 - m_2^*)] + D_\pm \Delta m_\pm \mp v_0 \nabla m_\pm - \nu m_\pm + \omega m_2 \quad ,
\end{aligned} \tag{S24}$$

and analogously for $z(x)$

$$\frac{\partial z}{\partial t} = \frac{z^*}{\tau_{\mathrm{z}}} \exp\left[\epsilon_{\mathrm{z}}(z - z^*)\right] - \exp\left[-\frac{\sigma - \sigma^*}{\sigma_{\mathrm{c}}}\right] z + D_z \Delta z \quad . \tag{S25}$$

Next, we introduce normalized concentration differences

$$\check{m}_\pm = \frac{m_\pm - m_{\pm^*}}{m_\pm^*} \tag{S26}$$

$$\check{m}_2 = \frac{m_2 - m_2^*}{m_2^*} \tag{S27}$$

$$\check{z} = \frac{z - z^*}{z^*} \quad , \tag{S28}$$

as well as normalized spatial inhomogeneity in mechanical tension

$$\check{\sigma} = \frac{\sigma - \sigma^*}{\sigma_c} = \frac{\sigma^*}{2\sigma_{\mathrm{c}}} \check{m} * (\chi_- - \chi_+) \quad . \tag{S29}$$

With these normalized quantities, the linearized dynamics reads

$$\frac{\partial \check{m}}{\partial t} = \frac{1}{\tau}\left(-\epsilon_\eta z^* \check{z} - \check{m}_\pm\right) - \frac{\check{m}_2}{\tau_2}\left(1 - \epsilon \frac{(\check{m}_2^*)^2}{m_\pm}\right) - \frac{m_2^*}{m_\pm^*}\frac{\check{m}_\pm}{\tau_2} \mp v_0 \nabla \check{m}_\pm + D_\pm \Delta \check{m}_\pm \quad , \tag{S30}$$

$$\frac{\partial \check{m}_2}{\partial t} = \frac{1}{\tau_2}\left(\check{m}_+ + \check{m}_- + 2\epsilon\, \check{m}_2 m_2^* - 2\check{m}_2\right) + D_m \Delta \check{m}_2 \quad , \tag{S31}$$

$$\frac{\partial \check{z}}{\partial t} = \frac{1}{\tau_z}\left(\epsilon_z \check{z} z^* + \check{\sigma} - \check{z}\right) + D_z \Delta \check{z} \quad . \tag{S32}$$

The dispersion relation was then computed with a computer algebra system, see Fig. S7A. From these dispersion relations, phase-diagrams as shown in Fig. 4 were computed.

For this analysis, we used $m_\pm^*$, $m_2^*$, $z^*$, $\sigma^*$, $\tau$, $\tau_2$, $\varepsilon_\eta$, $\varepsilon$, $\varepsilon_z$, $\nu_0$, $\sigma_c$ as effective parameters. To convert these effective parameters back to the parameters of the original model, we used the following relations

$$\eta^* = \frac{m_\pm^*}{\tau} \tag{S33}$$

$$\eta_0 = \eta^* \exp(\epsilon_\eta z^*) \tag{S34}$$

$$\beta_0 = \frac{1}{\tau} - \frac{2v_0}{l_a} \;\Rightarrow\; \beta = \tau^{-1} \tag{S35}$$

$$\omega^* = \frac{1}{\tau_2} = \beta_0 \exp(-\epsilon_\omega m_2^*) \;\Rightarrow\; \epsilon_\omega = -\frac{1}{m_2^*} \ln\left(\frac{\omega^*}{\beta_0}\right) \tag{S36}$$

$$\epsilon_\nu = \epsilon - \epsilon_\omega \tag{S37}$$

$$\nu^* = \omega^* \frac{m_2^*}{m_\pm^*} = \nu_0 \exp(\epsilon_\nu m_2^*) \;\Rightarrow\; \frac{1}{\nu_0} = \frac{1}{\nu^*} \exp(\epsilon_\nu m_2^*) \tag{S38}$$

$$\beta_z = \frac{1}{\tau_z} \exp(-\tau/\tau_c) \tag{S39}$$

$$\alpha_{z,0} = \beta_{z,0}\, z^* \exp(-\epsilon_z z^*) \tag{S40}$$

$$\alpha_z = \alpha_{z,0} \exp(\epsilon_z z) \quad . \tag{S41}$$

In addition, Fig. S7B shows the pattern length scale as function of the length of myosin and actin filaments.

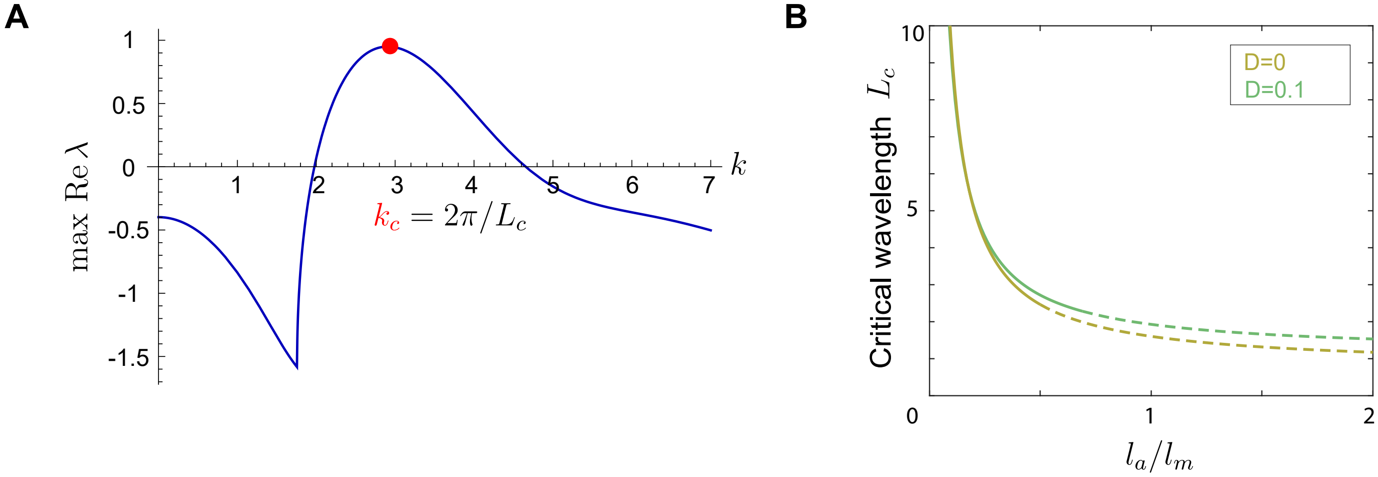


FIG. S7. **A.** Dispersion relation from linear stability analysis for model II of tension-driven myofibrillogenesis. For parameters, see Table S2. **B.** Critical wavelength $L_c$ as function of the ratio $l_a/l_m$ of actin and myosin filament lengths for two different values of the effective diffusion constant $D$.

*Model I with actin turn-over.* In the main text, we assumed for model I and II that actin filaments are static. For a modified model I, we include continuous turn-over of actin filaments (assuming immediate polymerization and depolymerization of actin filaments as in [20]), as well as preferential nucleation by Z-disc proteins. In a mean-field description, we thus obtain for the dynamics of the concentration fields $a_+(x)$ and $a_-(x)$ of actin filaments with minus-end pointing either in the positive or negative $x$-direction, respectively (see also Fig. 4) the dynamic equation

$$\frac{\partial a_\pm}{\partial dt} = -\beta_a a_\pm + \alpha_{a,0} + \alpha_{a,1}\ z * \chi_a\ , \tag{S42}$$

with depolymerization rate $\beta_a$, polymerization base rate $\alpha_{a,0}$, and a sur-plus polymerization rate $\alpha_{a,1}$ coupled to the local concentration of $z(x)$ of Z-disc proteins convoluted with a rectangular interaction function $\chi_a$ of amplitude $1/l_a$ and width $l_a$ centered at $\pm l_a/2$.

Results of an example computation are shown in Fig. S8, indicating an emergence of polarity-sorted actin patterns following the formation of the periodic pattern of Z-disc proteins.

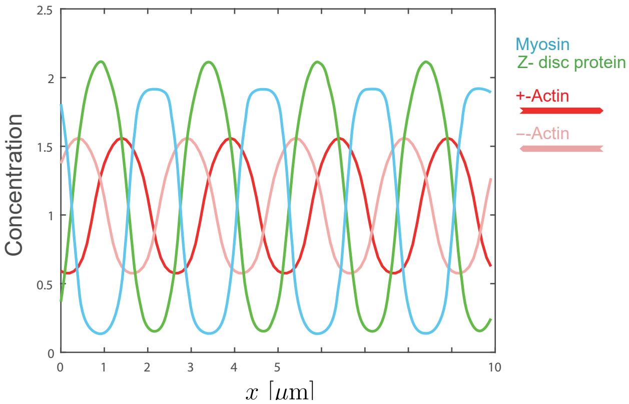


FIG. S8. Concentration profiles from mean-field simulation of a modified model I with additional actin turnover and preferential nucleation of actin by Z-disc proteins: myosin (blue), Z-disc proteins (green), actin filaments ('+'-polarity: red, '−'-polarity: rosé). Parameters: $l_a = 1$, $\alpha_{a,0} = 1$, $\alpha_{a,1} = 1$, $\beta_a = 2$; other parameters as in Table S1.

*Estimation of molecular force from surface tension.* We provide an order-of-magnitude estimate of the molecular force $f_0$ acting on individual actin crosslinkers as function of the macroscopic surface tension $\gamma$ as measured, e.g., for the actin cortex in [34]. By definition, a change $\Delta A$ in surface area of a thin film of crosslinked actin (with area $A$ and thickness $h$) requires a work $\Delta E = \gamma\,\Delta A$. If we idealize the actin meshwork as a cubic lattice of Hookean springs of equal length $a$ and spring constant $k$, the total elastic energy reads $E = \sum_i k\,x_i^2/2$, where $x_i$ is the strain of the $i$-th spring. The pre-strain $x = x_0$ of each spring sets the molecular force $f_0 = kx_0$ acting at each lattice node. Any additional strain $x = x_0 + \Delta x$ in $x$-direction increases this energy by $\Delta E = Nk\,x_0\Delta x$, while area increases by $\Delta A = A\Delta x/a$, where $N = Ah/a^3$ is the number of (relevant) springs. Hence, $f_0 = a^2\gamma/h$.
